# The single-cell atlas of the murine reproductive tissues during preterm labor

**DOI:** 10.1101/2022.04.27.489704

**Authors:** Valeria Garcia-Flores, Roberto Romero, Azam Peyvandipour, Jose Galaz, Errile Pusod, Bogdan Panaitescu, Derek Miller, Yi Xu, Li Tao, Zhenjie Liu, Adi L. Tarca, Roger Pique-Regi, Nardhy Gomez-Lopez

## Abstract

Preterm birth, the leading cause of perinatal morbidity and mortality worldwide, frequently results from the syndrome of preterm labor. Intra-amniotic infection is the best-established causal link to preterm labor, and involves premature activation of the parturition cascade in the reproductive tissues. Herein, we utilized single-cell RNA-sequencing (scRNA-seq) to generate a single-cell atlas of the murine uterus, decidua, and cervix in a model of infection-induced preterm labor. We show that preterm labor affects the transcriptomic profiles of specific immune and non-immune cell subsets. Shared and tissue-specific gene expression signatures were identified among affected cells. Importantly, determination of intercellular communications implicates specific cell types preterm labor-associated signaling pathways across tissues. Last, *in silico* comparison of murine and human uterine cell-cell interactions reveals conserved signaling pathways implicated in labor. Thus, scRNA-seq provides new insights into the preterm labor-driven cellular landscape and communications in the reproductive tissues.

## INTRODUCTION

Preterm birth is a devastating clinical condition that affects 15 million infants each year and is the leading cause of neonatal morbidity and mortality worldwide (Chawanpaiboon et al., 2019; Liu et al., 2015b). Spontaneous preterm birth often results from preterm labor, a syndrome for which multiple etiologies have been proposed (Goldenberg et al., 2008; Romero et al., 2014a). Among these, the best-established causal link to preterm birth is intra-amniotic infection, a clinical condition resulting from the invasion of microbes into the amniotic cavity (Agrawal and Hirsch, 2012; Bastek et al., 2011; Combs et al., 2014; Gibbs et al., 1992; Gomez et al., 1995; Gonçalves et al., 2002; Keelan et al., 2003; Romero et al., 2019; Theis et al., 2020). The most frequently detected bacteria in amniotic fluid of women diagnosed with intra-amniotic infection include genital mycoplasmas, *Streptococcus agalactiae*, *Gardnerella vaginalis*, and *Escherichia coli*, among others (Burnham et al., 2020; DiGiulio et al., 2010; Gibbs et al., 1982; Gravett et al., 1986; Oh et al., 2019; Romero et al., 2015a; Romero et al., 2014b; Romero et al., 2015b; Romero et al., 2015c; Romero et al., 1989; Yoon et al., 1998; Yoon et al., 2019). Human descriptive studies have consistently shown that such microbial invasion of the amniotic cavity is accompanied by a local acute inflammatory response that includes the infiltration of leukocytes into the amniotic cavity (inc. amniotic fluid (Galaz et al., 2020a; Galaz et al., 2020b; Gomez-Lopez et al., 2019a; Gomez-Lopez et al., 2019b; Gomez-Lopez et al., 2021b; Gomez-Lopez et al., 2017b; Gomez-Lopez et al., 2019c; Gomez-Lopez et al., 2018b; Gomez et al., 1994; Martinez-Varea et al., 2017; Romero et al., 1991; Romero et al., 1993a; Romero et al., 1993b; Yoon et al., 1996) and placental tissues (Gomez-Lopez et al., 2017a; Guzick and Winn, 1985; Hillier et al., 1988; Kim et al., 2015a; Kim et al., 2009; Kim et al., 2015b; Odibo et al., 1999; Oh et al., 2021; Oh et al., 2011; Olding, 1970; Pacora et al., 2002; Park et al., 2009; Redline, 2012; Redline et al., 2003; Svensson et al., 1986; van Hoeven et al., 1996; Yoon et al., 2001; Zhang et al., 1985)) as well as the reproductive tissues (Keski-Nisula et al., 2003; Makieva et al., 2017). More recently, animal models coupled with omics technologies have been utilized to strengthen this concept and establish causality between intra-amniotic infection and the inflammatory milieu observed in the reproductive tissues (e.g., uterus, decidua, and cervix) that serve to orchestrate the premature activation of the common pathway of labor (Migale et al., 2016; Motomura et al., 2020a; Toothaker et al., 2020; Willcockson et al., 2018). However, the simultaneous investigation of the cellular landscape and interaction networks at single-cell resolution in the reproductive tissues implicated in preterm parturition has not been undertaken.

Single-cell technology has emerged as a useful tool for evaluating cellular composition, transcriptomic activity, and communication networks in gestational and reproductive tissues (Huang et al., 2021; Nelson et al., 2016; Pavličev et al., 2017; Pique-Regi et al., 2019; Suryawanshi et al., 2018; Tsang et al., 2017; Vento-Tormo et al., 2018). Indeed, we have applied single-cell RNA-sequencing (scRNA-seq) to investigate the physiological and pathological processes of labor in the placenta and extraplacental membranes (Pique-Regi et al., 2019). More recently, we utilized scRNA-seq to unravel the myometrial cell types that participate in the normal process of term parturition as well as key cell-cell interactions taking place in this compartment (Pique-Regi et al., 2022). Importantly, the discovery of single-cell signatures derived from the placental tissues and myometrium possesses translational value as these can serve as potential non-invasive biomarkers of labor progression and/or obstetrical disease (Gomez-Lopez et al., 2022; Pique-Regi et al., 2022; Pique-Regi et al., 2019; Tarca et al., 2021; Tarca et al., 2019; Tsang et al., 2017).

In the current study, we utilized scRNA-seq coupled with an allogeneic murine model of intra-amniotic infection to investigate the cellular landscape and cell-cell communications in the reproductive tissues (uterus, decidua, and cervix) during the process of preterm labor. We utilized a murine model of preterm labor and birth induced by the intra-amniotic inoculation of *E. coli*, and assessed cervical shortening to establish the timing of active preterm labor. Next, using scRNA-seq and computational approaches, we generated a single-cell atlas of the uterus, decidua, and cervix during preterm labor as well as their cell type-specific transcriptomic activity. In addition, we established the cell-cell communication networks between cell types in each tissue during preterm labor and identified key signaling pathways implicated in this process. Last, we integrated cell-cell signaling pathways derived from the murine uterus with those from the human myometrium during the processes of preterm and term labor, respectively, to demonstrate conserved labor-associated signaling.

## RESULTS

### A single-cell atlas of the murine reproductive tissues during preterm labor induced by *E. coli*

Intra-amniotic infection has been documented as inducing inflammatory changes in the tissues surrounding the amniotic cavity (Faro et al., 2019; Motomura et al., 2021; Motomura et al., 2020b; Senthamaraikannan et al., 2016). To investigate such changes at single-cell resolution, we first established a murine model of preterm labor and birth induced by the intra-amniotic inoculation with *E. coli*, one of the microorganisms commonly identified in the amniotic fluid of women with intra-amniotic infection (Gibbs et al., 1982; Romero et al., 2015c; Romero et al., 1989; Yoneda et al., 2016a). Mice with an allogeneic pregnancy underwent ultrasound-guided intra-amniotic injection of *E. coli* or vehicle control on 16.5 days *post coitum* (dpc) (Figure 1A). Intra-amniotic inoculation with *E. coli* reduced gestational length in the majority of dams (Figure 1B), resulting in an 83.3% (5/6) rate of preterm birth (Figure 1C). We then intra-amniotically injected a second cohort of mice with *E. coli* or PBS to perform tissue collection for single-cell analyses. To ensure that the *E. coli-*injected mice were undergoing preterm labor at the time of tissue collection, we utilized ultrasound to evaluate cervical length just prior to intra-amniotic injection and again 24 h later as a readout of cervical effacement (Figure 1D). Cervical shortening was observed in all dams that received intra-amniotic *E. coli* at 24 h post-injection, indicating that these dams were in active preterm labor at the time of tissue collection, whereas no cervical shortening was observed in controls (Figure 1E). Therefore, intra-amniotic inoculation with *E. coli* represents a translational model that resembles the clinical scenario of intra-amniotic infection leading to preterm labor and birth.

**Figure 1.**
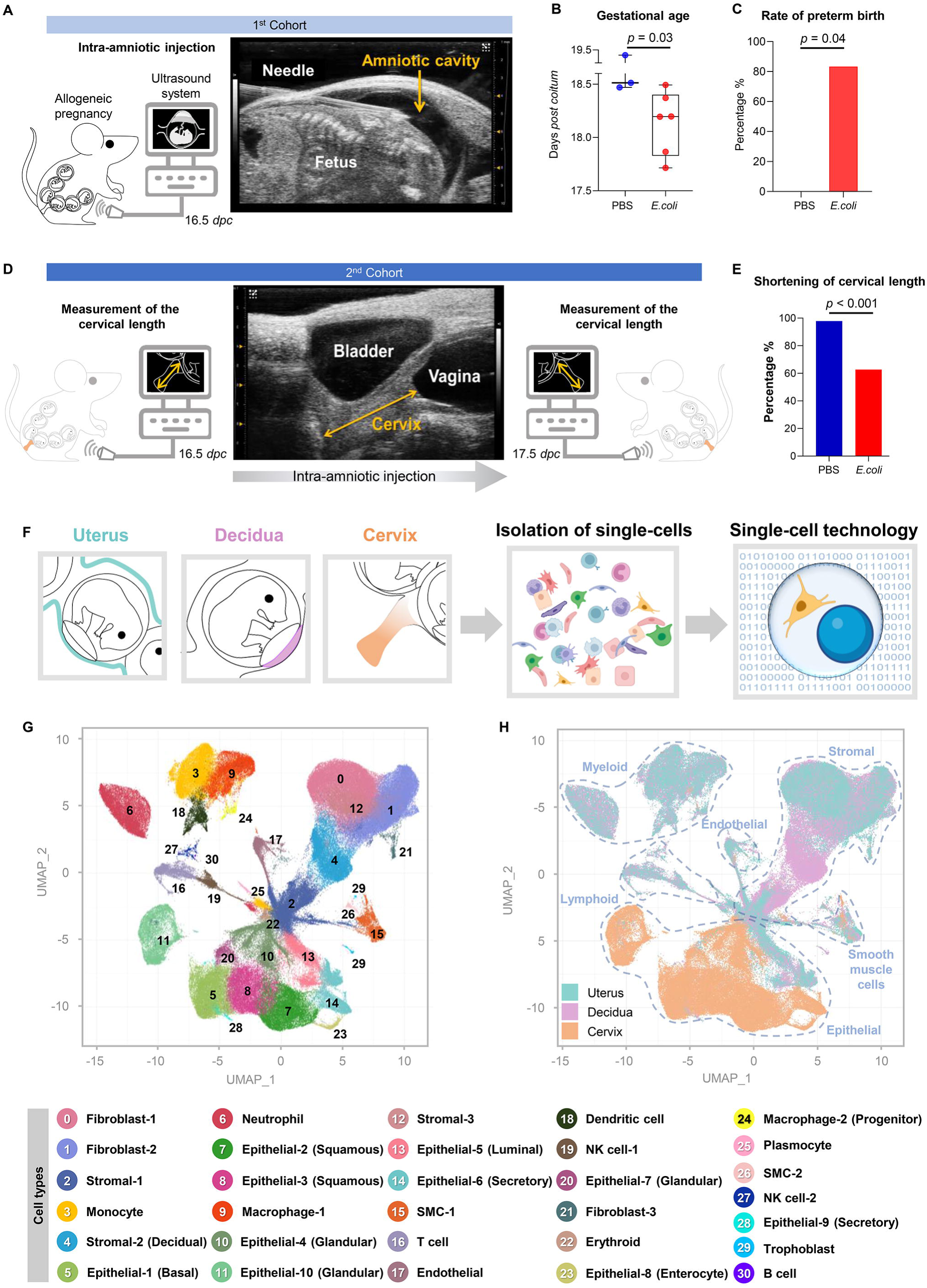
Single-cell atlas of the murine reproductive tissues during preterm labor induced by intra-amniotic infection. **(A)** Experimental design for the ultrasound-guided intra-amniotic injection of *E. coli* or PBS (vehicle control) into pregnant mice on 16.5 days *post coitum* (dpc) (n = 3-6 per group). Mice were monitored to determine pregnancy outcomes **(B-C)**. Gestational age and preterm birth rate of dams intra-amniotically injected with *E. coli or* PBS. Gestational age was compared using a 2-sided Mann-Whitney U-test, and preterm birth rates were compared using a 2-sided Fisher’s test. P < 0.05 was considered significant. **(D)** Experimental design for the determination of cervical length on 16.5 dpc, prior to intra-amniotic injection of *E. coli* or PBS, and 24 h later (17.5 dpc) (n = 6-9 per group). **(E)** Cervical length of dams intra-amniotically injected with *E. coli* or PBS at 16.5 and 17.5 dpc. Cervical length was compared between time points using a 2-sided Mann-Whitney U-tests. P < 0.05 was considered significant. The change in cervical shortening was calculated by considering the measurement at 16.5 dpc as 100%. **(F)** Diagram illustrating the generation of single-cell suspensions from the uterus, decidua, and cervix collected for single-cell RNA-sequencing experiments (scRNA-seq) (n = 4 per group). **(G)** Uniform manifold approximation and projection (UMAP) plot showing all cell types present in the uterus, decidua, and cervix. **(H)** UMAP color-coded plot showing tissue-specific predominance of distinct cell types in the uterus (blue), decidua (pink), and cervix (orange). Blue dotted lines distinguish major cell types: myeloid, endothelial, stromal, smooth muscle, epithelial, and lymphoid. Abbreviations used: SMC, smooth muscle cell; NK cell, natural killer cell.

Preterm parturition includes the activation of the common pathway of labor that comprises increased uterine contractility, the triggering of local immune response in the decidual tissues, and cervical dilatation (Lopez Bernal, 2003; Norwitz et al., 1999; Romero et al., 2014a; Romero et al., 2006; Romero et al., 1994; Smith, 2007). Therefore, to establish a single-cell atlas of the murine reproductive tissues in preterm labor, we utilized the uterus, decidua, and cervix of dams that received intra-amniotic inoculation with *E. coli* in the active phase of parturition (17.5 dpc) for single-cell RNA-sequencing (scRNA-seq) (Figure 1F). We identified 31 cell clusters across the uterus, decidua, and cervix that corresponded to multiple cell types: smooth muscle (2 clusters), epithelial (10 clusters), fibroblast (3 clusters), stromal (3 clusters), Endothelial, Neutrophil, Monocyte, macrophage (2 clusters), Dendritic Cell, T cell, B cell, NK cell (2 clusters), Erythroid, Plasmocyte, and Trophoblast (Figure 1G). The heterogeneous and distinct cellular composition of the uterus, decidua, and cervix was highlighted by assigning tissue identity to each cell cluster (Figure 1H). In control dams, the uterus, decidua, and cervix each displayed a distinct basal cellular repertoire: the uterus showed a predominance of fibroblast (clusters 0 and 1) and non-decidual stromal (clusters 2 and 12) cell types, and the decidua also included a unique subset of stromal cells (cluster 4) (likely corresponding to conventional decidual stromal cells) (Figure S1A). The uterus and decidua of control mice also included modest populations of innate immune cells such as Monocyte and macrophage subsets as well as lymphocytes such as T cell, NK cell-1, NK cell-2, and B cell (Figure S1A), likely representing the resident immune populations that have been characterized in human and murine tissues (Bartmann et al., 2014; Li et al., 2018; Pique-Regi et al., 2022; Pique-Regi et al., 2019; Trundley and Moffett, 2004; Vento-Tormo et al., 2018). By contrast with the uterus and decidua, the cervix of control mice comprised a diverse compartment of epithelial subsets (clusters 5, 7, 8, 10, 11, 14) and other major cell types (Figure S1A), as previously shown (Chumduri et al., 2021; Koh et al., 2019; Zhao et al., 2021). Immune cells were scarce in the cervix, although a modest Macrophage-1 population was observed (Figure S1A) that is consistent with prior reports of cervical cell composition in late gestation (Dobyns et al., 2015; Dubicke et al., 2016; Osman et al., 2003; Timmons et al., 2009). Together, these data provide an overview of the single-cell composition and diversity in the murine uterus, decidua, and cervix in late murine pregnancy. Moreover, by dissecting the cellular repertoire of the cervix we demonstrate the underappreciated heterogeneity of this compartment.

### Preterm labor induced by *E. coli* dysregulates the repertoire of immune and non-immune cell types in the reproductive tissues

We then examined the effects of preterm labor on the abundance of each cell type identified across all tissues (Figure 2A) as well as within the uterus, decidua, and cervix (Figure 2B-D, Figure S1B, and Table S1). During preterm labor, a relative increase in innate immune cell clusters such as Monocytes, macrophages, Dendritic cells, and Neutrophils (clusters 3, 6, 9, and 18) was observed in the uterus, decidua, and, to a lesser extent, the cervix (Figure 2B-D and Figure S1B). The NK cell-2 and Plasmocyte subsets in the uterus and decidua also showed changes with preterm labor (Figure 2B-C and Figure S1B). Moreover, Dendritic cells were increased in the decidua (Figure 2C and Figure S1B), with a similar tendency observed in the uterus (Figure 2B and Figure S1B). Interestingly, the Macrophage-1 cell type was decreased in the uterus with preterm labor (Figure 2B and Figure S1B). Notably, the T-cell population (cluster 16) also appeared to increase in the uterus and decidua with preterm labor (Figure 2B&C and Figure S1B), which is consistent with prior studies implicating T-cell infiltration and activation as a component of parturition (Arenas-Hernandez et al., 2019; Gomez-Lopez et al., 2009; Gomez-Lopez et al., 2011a; Gomez-Lopez et al., 2011b; Gomez-Lopez et al., 2013; Sindram-Trujillo et al., 2003; Slutsky et al., 2019; Vargas et al., 1993). Although not visually apparent from the UMAP plots (Figure S1B), both the uterus and decidua showed a substantial decrease in non-immune subsets such as Fibroblast-1, -2, and Stromal-3 with preterm labor (Figure 2B&C), with Stromal-2 also showing modest changes in the uterus (Figure 2B). Interestingly, a subset of epithelial cells (cluster 11, Epithelial-10) that was largely absent in the uterus and decidua of controls became apparent in preterm labor (Figure S1B), suggesting labor-induced differentiation or activation of these cells. By contrast with the uterus and decidua, the cervix only showed changes in two cell types: Neutrophil and Epithelial-8 were both increased with preterm labor (Figure 2D), indicating that a modest cellular response to intra-amniotic infection occurs in this tissue. We also evaluated whether cells of fetal origin were represented among the populations of the uterus, decidua, and cervix during preterm labor (Figure S1C-E). A small population of fetal cells (Trophoblasts) was detected in the uterus and decidua, which is consistent with prior single-cell studies of the human myometrium (Pique-Regi et al., 2022) and may represent residual placental cells attached to the uterus and decidua.

**Figure 2.**
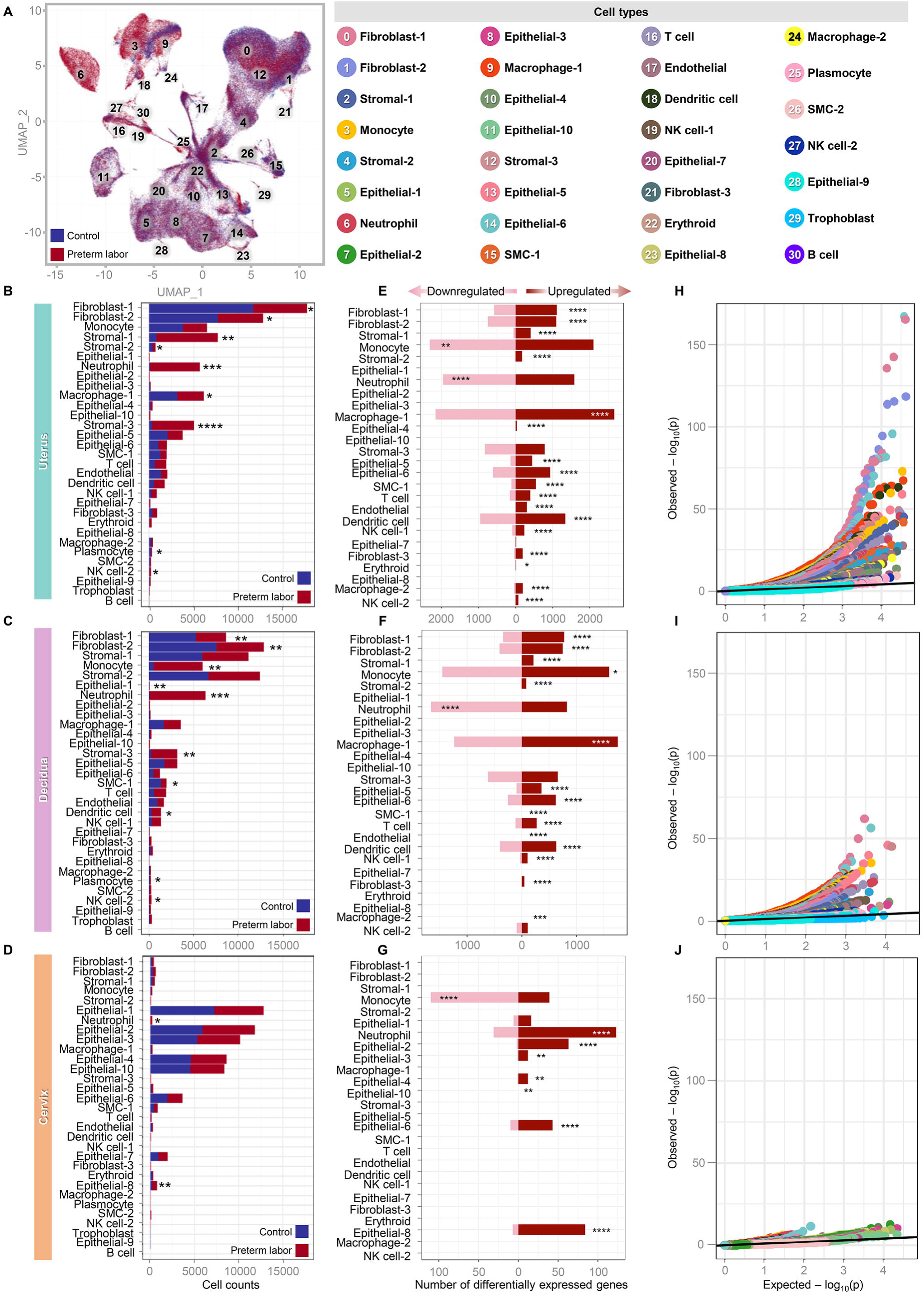
Preterm labor induced by *E. coli* dysregulates the repertoire and gene expression of immune and non-immune cell types in the reproductive tissues. **(A)** Color-coded uniform manifold approximation and projection (UMAP) plot showing the effects of preterm labor on the abundance of specific cell types (shown in red) compared to control (shown in blue). **(B-D)** Bar plots showing the numbers of each cell type in the uterus, decidua, and cervix. The comparison of cell numbers between the two study groups for each cell type was performed using a 2-sided t-test. * p < 0.05, ** p < 0.01, *** p < 0.001, **** p < 0.0001. **(E-G)** Bar plots showing the numbers of differentially expressed genes (DEGs) induced by preterm labor in cell type in the uterus, decidua, and cervix. Red and pink bars indicate upregulated and downregulated DEGs, respectively (derived from DESeq2, q < 0.1). The comparisons of the fraction of downregulated and upregulated DEGs in each cell type between the study groups were calculated using two-sided binomial tests. * q < 0.05, ** q < 0.01, *** q < 0.001, **** q < 0.0001. **(H-J)** Quantile-quantile plot showing differential expression of genes analyzed for selected enriched cell types from the uterus, decidua and cervix. Deviation above the 1:1 line (solid black line) indicates enrichment. Abbreviations used: SMC, smooth muscle cell; NK cell, natural killer cell.

To validate the leukocyte infiltration of the uterus, decidua, and cervix indicated by our single-cell data, we undertook a series of histological and immunohistochemistry analyses (Figure S2). We observed collagen degradation in the uterine and cervical tissues with preterm labor, and mucin production by cervical cells appeared to increase compared to control tissues (Figure S2A-C). Histological changes in preterm labor were accompanied by increased CD45+ leukocyte infiltration in the uterus and decidua (Figure S2D-F). Uterine leukocytes were more evenly distributed among neutrophils, monocytes, and macrophages, whereas decidual leukocytes were predominantly neutrophils and, to a lesser extent, monocytes (Figure S2G-I). Similar to our scRNA-seq results, the leukocyte abundance in the cervical tissues was largely comparable between the control and preterm labor groups (Figure S2I).

Taken together, our scRNA-seq indicate a shift in the cellular composition of the murine uterus, decidua, and cervix that accompanies preterm labor.

### Preterm labor induced by *E. coli* dysregulates the gene expression of immune and non-immune cell types in the reproductive tissues

Given that preterm labor altered the cellular composition of the uterus, decidua, and cervix, we next explored whether this inflammatory process also resulted in transcriptomic changes to the identified cell types. Consistent with their altered abundance, multiple fibroblast, stromal, and epithelial cell types in the uterus and decidua displayed upregulated gene expression with preterm labor (Figure 2E&F and Table S2), while the cervical non-immune cells with upregulated gene expression were exclusively epithelial (Figure 2G and Table S2). Interestingly, innate immune cell types showed strong dysregulation of gene expression in both directions that was inconsistent among tissues: while Monocyte showed more downregulated differentially expressed genes (DEGs) in the uterus (Figure 2E) and cervix (Figure 2G), this cell type showed more upregulated DEGs in the decidua (Figure 2F). Neutrophil showed stronger downregulation of DEGs in the uterus and decidua (Figure 2E&F), whereas in the cervix DEGs were primarily upregulated in this cell type (Figure 2G). Macrophage-1, Dendritic cell, and NK cell-1 consistently displayed predominantly upregulated DEGs in the uterus and decidua (Figure 2E&F), and were not represented in the cervix as previously noted. In addition, the uterine Macrophage-2 and NK cell-2 populations displayed upregulated DEGs (Figure 2E), which was not observed in other tissues (Figure 2F&G). Although not as abundant as innate immune cells, the T cell population also displayed upregulated DEGs with preterm labor in the uterus and decidua (Figure 2E&F). Moreover, quantile-quantile plots of DEGs from enriched cell types indicate that the uterus is the tissue most affected by the process of labor (Figure 2H-J). Thus, preterm labor primarily induces gene expression in the dominant cell types from each tissue; yet, the substantial amount of downregulated gene expression in innate immune cells may indicate an immunological switch from one transcriptomic program to another to combat infection.

### Preterm labor induced by *E. coli* involves conserved cell types that display distinct processes in the reproductive tissues

The transcriptomic profiling of cell types suggested that specific subsets show conserved responses with preterm labor across the reproductive tissues. Therefore, we next focused on the shared preterm labor-specific gene expression among the uterus, decidua, and cervix. The Venn diagram displayed in Figure 3A highlights the overlap in DEGs across tissues, particularly the uterus and decidua. Correlation analyses indicated stronger relationships between preterm labor and gene expression changes in the uterus and decidua than in the cervix (Figure S3A), which was reflective of the total preterm labor-associated DEGs in each tissue. This observation was further confirmed by the correlation between the gene expression profiles of the uterus and decidua, which was stronger than the correlations between the decidua and cervix or the uterus and cervix (Figure S3B). Given that the uterus, decidua, and cervix all displayed some degree of correlation for preterm labor-associated gene expression, we evaluated the cell type-specific transcriptomic changes that were conserved across all three tissues. We found that innate immune cell types (Monocyte and Neutrophil) as well as non-immune cell types (Epithelial-3, -4, -6 and Endothelial) showed conserved gene expression changes associated with preterm labor across the uterus, decidua, and cervix (Figure 3B). We reasoned that, although the transcriptome profiles of specific cell types were affected across all tissues, such cells may display distinct biological processes according to their location. Gene Ontology (GO) analysis of the Neutrophil cell type in the uterus, decidua, and cervix revealed that, while these cells shared some processes such as “response to bacterium” and “response to lipopolysaccharide”, processes specific to Neutrophils in each tissue were also observed (Figure 3C). Uterine Neutrophils showed enrichment of processes related to cytokine signaling and anti-viral response, whereas decidual Neutrophils showed enrichment of cellular activation-associated processes (Figure 3C). In the cervix, unique enriched Neutrophil processes were primarily associated with response to external stimuli and bacteria (Figure 3C). Uterine and decidual Monocyte and Macrophage-1 cells also shared enriched processes related to cytokine production and response to bacteria/lipopolysaccharide, with decidual Monocytes also showing enrichment of activation-associated processes (Figure 3D). By contrast, cervical Monocytes displayed highly distinct processes related to protein synthesis and humoral immune response (Figure 3D), suggesting that such cells may functionally differ from their counterparts in the uterus and decidua. Epithelial-6, which had sufficient DEGs to perform GO analysis in all three tissues, displayed largely consistent processes across the uterus, decidua, and cervix that were related to inflammation, anti-bacterial response, and cytokine production (Figure 3E). The uterine Epithelial-4 cell type displayed enrichment of several chemotaxis-associated processes, suggesting involvement in leukocyte recruitment to this tissue, whereas the cervical Epithelial-4 showed enrichment of effector functions such as production of NO, IL-1, and IFNγ (Figure 3E). Epithelial-3, which only displayed sufficient DEGs for GO analysis in the cervix, showed enrichment of multiple processes related to promotion of B-cell and antibody responses (Figure 3E). Thus, the conserved cell types affected by preterm labor in the uterus, decidua, and cervix each display distinct enrichment of biological processes, suggesting that similar cell types display tissue-specific functions in the context of intra-amniotic infection leading to preterm labor. Together with the observed increase in cervical Epithelial-8 cell counts with preterm labor, it is possible that the upregulation of inflammatory gene expression represents an infection-induced differentiation of cervical epithelial cells to better participate in host defense mechanisms in this compartment.

**Figure 3.**
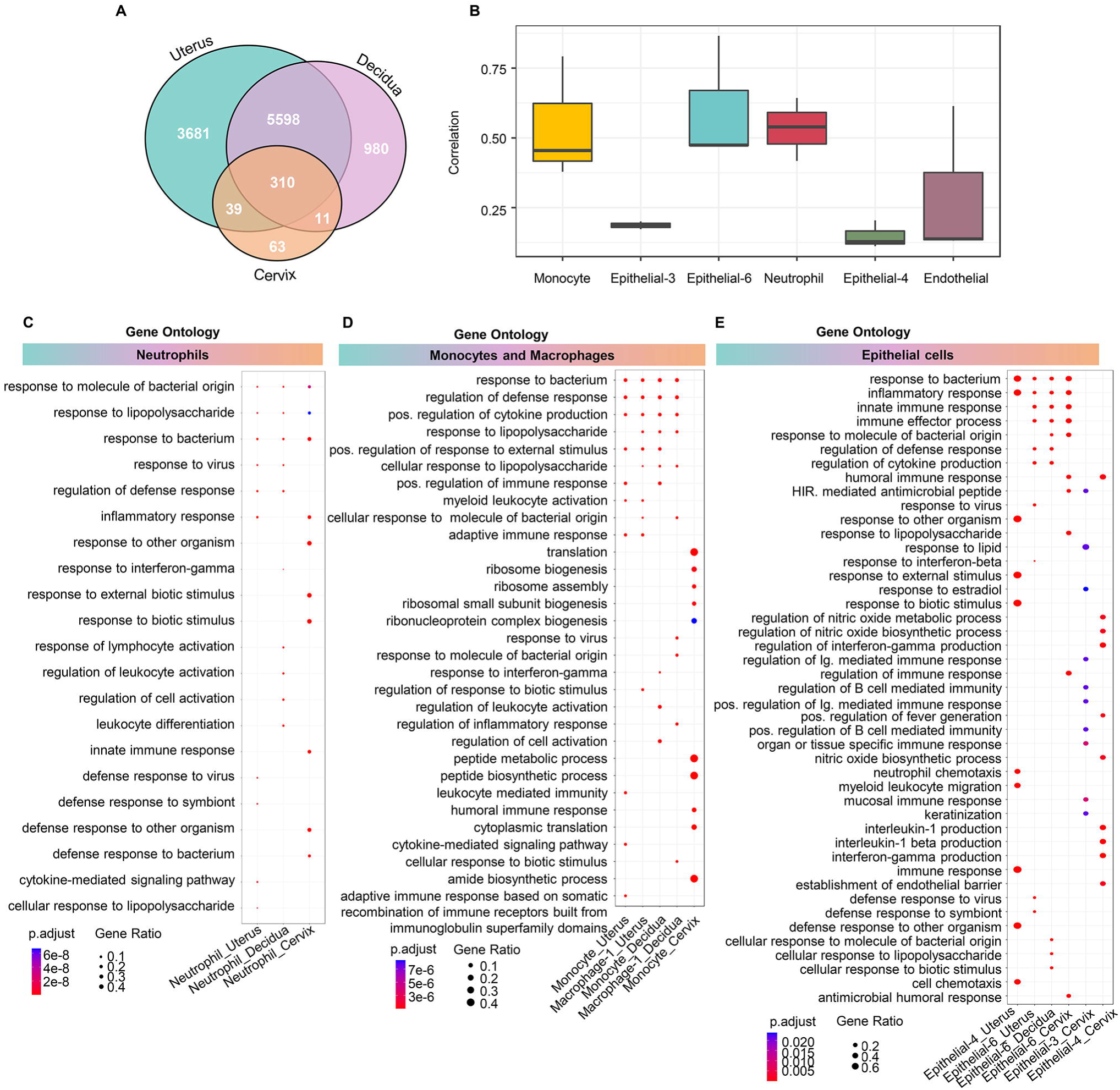
Preterm labor induced by *E. coli* involves conserved cell types that display distinct processes in the reproductive tissues. **(A)** Venn diagrams showing the numbers of differential expressed genes (DEGs, q < 0.1) that are unique to or shared among the uterus, decidua, and cervix. **(B)** Box plots showing the correlation of specific cell types affected by preterm labor and conserved across the uterus, decidua, and cervix using the Spearman’s method. Cluster profiler dot plots showing the preterm labor-associated Gene Ontology (GO) biological processes that are unique or shared among **(C)** Neutrophil, **(D)** Monocyte and Macrophage, and (E) Epithelial cell types from the uterus, decidua, and cervix. The size and color of each dot represent gene ratio and significance level, respectively. 1-sided Fisher’s exact tests were used. Abbreviations used: SMC, smooth muscle cell; NK cell, natural killer cell.

To further infer cellular functionality in preterm labor, we then utilized the Kyoto Encyclopedia of Genes and Genomes database to evaluate the pathways enriched in labor-associated DEGs in each cell type (Figure S3C). Strikingly, both immune and non-immune cell types with altered gene expression in preterm labor showed enrichment of immunological pathways such as “cytokine-cytokine receptor interaction”, “NOD-like receptor signaling pathway”, and “viral protein interaction with cytokine and its receptor” across the three tissues (Figure S3C). Such findings are consistent with previous studies showing the upregulation of immune-related pathways in decidual endothelial (Huang et al., 2020) and stromal cells (Huang et al., 2021) from women with labor. Additional inflammatory pathways such as “NF-kappa B signaling pathway” and “Toll-like receptor signaling pathway” were also represented, to a lesser extent, by immune cells (e.g., NK cells and Neutrophil) as well as non-immune cells such as epithelial cells, including in the cervix (Figure S3C).

We also investigated the biological processes enriched in several non-immune cell subsets that were conserved between the uterus and decidua with preterm labor (Figure S4). Interestingly, Stromal-1 and Stromal-2 in the uterus showed largely similar enrichment of biological processes, as did the Stromal-1 and Stromal-2 cell types in the decidua (Figure S4A). Yet, these cell types differed between tissues, given that the decidual stromal cells were enriched for leukocyte migration and chemotaxis whereas the uterine stromal cells showed enrichment for response to immune signaling (Figure S4A). The Fibroblast-1 and Stromal-3 cell types showed less diversity in their enriched processes compared between the uterus and decidua, with the former being associated with host defense against infection and the latter associated with immune activation, including adaptive immunity (Figure S4B). Fibroblast-2 and Fibroblast-3 were also comparable between the uterus and decidua; however, the decidual Fibroblast-3 showed more striking enrichment of responses to microorganisms and cytokine signaling (Figure S4C). Finally, the uterine and decidual Endothelial cell types displayed similar enrichment of processes related to host defense, innate immunity, and cytokine signaling, with the decidual subset showing modestly higher enrichment for processes related to neutrophil migration (Figure S4D).

Taken together, these data indicate that the uterus, decidua, and cervix contain cell types that display distinct tissue-specific gene expression profiles in preterm labor, pointing to differing functional roles for these cells in the host response to intra-amniotic infection. Yet, there is an overall tendency for the enrichment of similar immunological pathways in both immune and non-immune cell types across tissues, likely as part of the common host response to intra-amniotic infection.

### Preterm labor influences cell-cell communications in the reproductive tissues

Having established that preterm labor drives distinct transcriptomic changes in specific cell types in the uterus, decidua, and cervix, we next leveraged our single-cell data to elucidate cell-cell communication networks in these tissues.

#### Cell-cell communications in the uterus

The uterus is a highly heterogeneous organ with multiple described regions that differ in cellular composition and function (Bukowski et al., 2006; Danforth and Ivy, 1949; Liu et al., 2015a; Mosher et al., 2013; Patwardhan et al., 2015; Pollard et al., 2000; Schwalm and Dubrauszky, 1966; Sooranna et al., 2006; Sparey et al., 1999; Wikland et al., 1982). To unravel the intercellular communications taking place in the murine uterus with preterm labor, we performed a correlation analysis across preterm labor-associated genes for each pair of identified cell types (Figure S5A). The strongest correlations were observed for non-immune cell types, such as stromal, epithelial, fibroblast, smooth muscle, and Endothelial cell types (Figure S5A), suggesting that these cells exhibit similar changes in gene expression with preterm labor. Innate and adaptive immune cell types also showed moderate correlations; namely, T cell, NK-cell-1, NK-cell-2, Macrophage-1, Macrophage-2, Monocyte, and Dendritic cell (Figure S5A). The cell types with the weakest correlations were primarily epithelial subsets as well as Neutrophil, Erythroid, Stromal-3, and SMC-2 (Figure S5A), indicating that such cells show more distinct gene expression changes with preterm labor.

We next used CellChat to infer cell-cell communications within the uterus using our single-cell gene expression data and a database of established interactions between signaling ligands, receptors, and their cofactors (Jin et al., 2021). Signaling pathways that were enriched or diminished in preterm labor, as well as those that were unaffected by this process, are shown in Figure S5B and Table S3. The alluvial plots shown in Figure 4A&B display the major cell-cell communication processes taking place in the uterus as well as the cell types that participate as senders or receivers in each process. Both innate and adaptive immune cell subsets (Neutrophil, Macrophage-1, Monocyte, Dendritic cell, NK cell-1, T-cell) contribute to the top signaling pathways implicated in preterm labor, such as CCL, CXCL, complement, IFN-I and IFN-II, IL-1, and IL-6 (Figure 4A&B). Notably, multiple non-immune subsets also participate in these processes; namely, fibroblast, stromal, epithelial, SMC, and Endothelial cell types (Figure 4A&B). While both immune and non-immune cell types served as receivers of preterm labor-associated signaling, specific responders to each pathway could be distinguished (Figure 4B). For example, the signaling pathway of IL-6, which is commonly utilized as a biomarker of intra-amniotic inflammation (Yoon et al., 2001), was primarily driven by immune cell types (Figure 4A); yet, the receiver cells for this pathway were non-immune subsets (Figure 4B). Conversely, the primary senders for the Annexin signaling pathway were non-immune cell types (Figure 4A), with the downstream receivers being predominantly immune cells (Figure 4B). The changes in cell-cell communication occurring as a result of preterm labor were visualized using the arrow plot in Figure 4C, where the directionality of each cell type arrow reflects the propensity for increased outgoing and/or incoming interaction strength. Cell types such as Macrophage-2, Stromal-2, Stromal-3, and Fibroblast-3 showed primarily even increases in incoming and outgoing signaling with preterm labor (Figure 4C). Other cell types were more biased towards incoming interactions, such as Macrophage-1, Neutrophil, Dendritic cell, Plasmocyte, Monocyte, and Epithelial-6, or outgoing interactions, such as Stromal-1, Fibroblast-1, and Fibroblast-2 (Figure 4C). Interestingly, several cell types showed a net decrease in signaling with preterm labor; namely, T cell and SMC-1 (Figure 4C).

**Figure 4.**
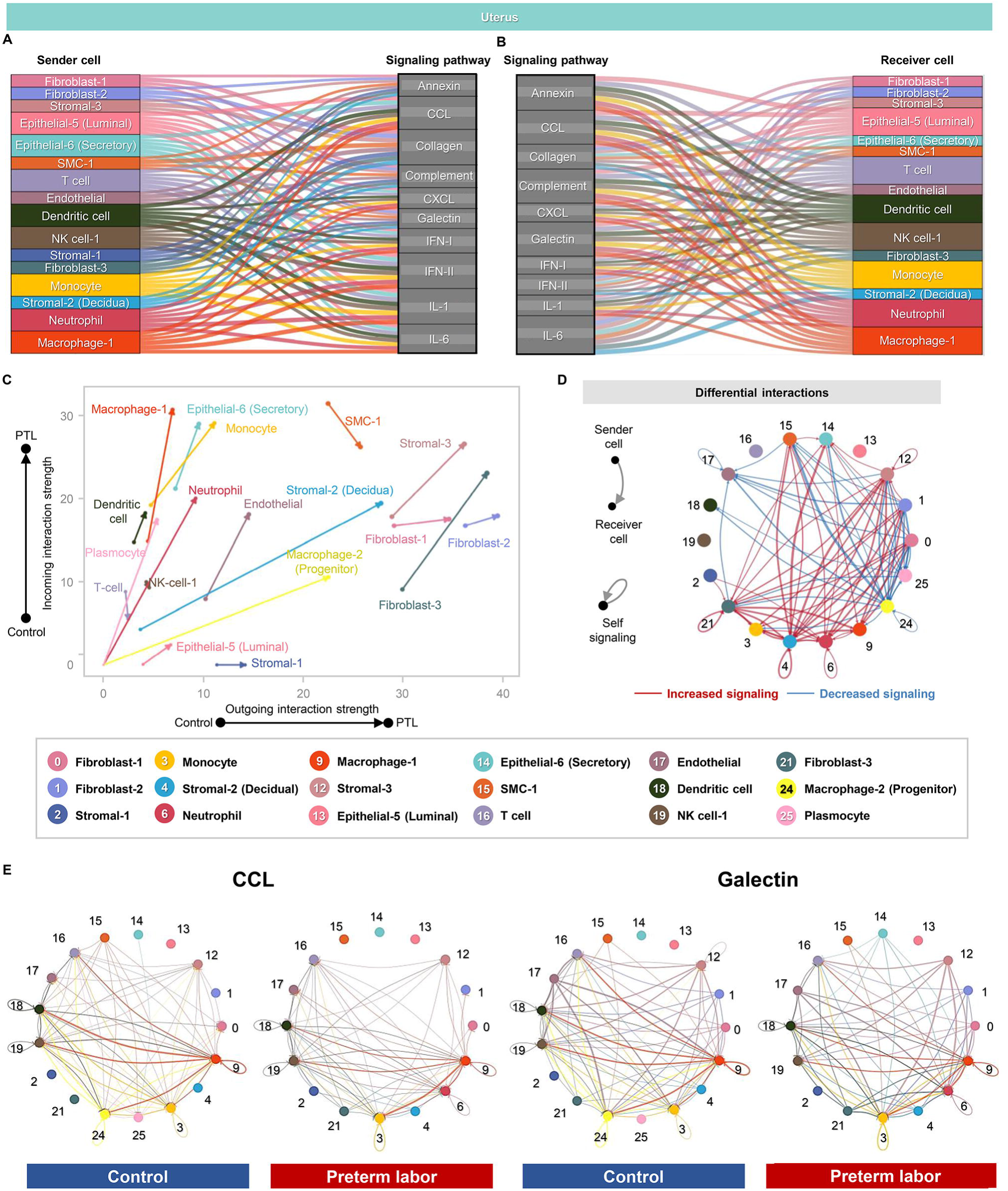
Cellular interactions in the uterus during preterm labor. **(A-B)** Alluvial plots showing the roles of specific cell types as senders or receivers of preterm labor-associated signaling in the uterus based on selected enriched signaling pathways comparing the overall information flow within networks between preterm labor and control derived from CellChat (full list of pathways in Figure S5B). Connecting lines are color-coded and represent the participation of each cell type as senders or receivers of the indicated signaling pathway. Line thickness is proportional to the strength of interaction. **(C)** Arrow plot showing changes in outgoing and incoming interactions strength between preterm labor (point of the arrow) and control condition (base of the arrow) for specific cell types in the uterus. **(D)** Circle plots showing the top 25% of aggregated interactions among cell types in the uterus for control and preterm labor groups. Each node represents a cell type and the interaction is shown by lines color-coded based on the sender cell. **(E)** Circle plots showing the top 25% increased (red) or decreased (blue) signaling interactions in the uterus for specific pathways in preterm labor compared to controls.

The top 25% of aggregated cellular interactions in the uterus were then contrasted between the control and preterm labor groups, emphasizing the overall increase in cell-cell signaling with preterm labor as well as the incorporation of new signaling pathways from cell types that were rarely present in control tissues, such as Neutrophil (Figure 4D). Interestingly, while Macrophage-1 signaling was increased, Macrophage-2 signaling decreased, which could indicate a homeostatic role for the latter subset that is diminished in preterm labor, as previously reported (Gomez-Lopez et al., 2021a). Next, we examined the top contributors within uterine cell-cell signaling pathways enriched with preterm labor (Figure 4E). We found that macrophage subsets and Dendritic cells were primary contributors to CCL signaling between control uterine cell types, and such signaling was further strengthened in preterm labor (Figure 4E). By contrast, the Galectin signaling pathway, already enriched in control uterine tissues, was upregulated in new cell types in preterm labor (e.g., Epithelial-6) and diminished in others (e.g., Macrophage-2) (Figure 4E).

We also explored the changes in cell type-specific expression of genes related to progesterone and prostaglandin signaling in the uterus (Figure S5C and Figure S6A). As expected, progesterone-related gene expression was consistently downregulated across uterine cell types in preterm labor (Figure S5C). Prostaglandin-related gene expression showed more activity in the uterus than in other tissues (Figure S6); yet, preterm labor-associated changes in each gene were consistent across uterine cell types (Figure S6A), further supporting the involvement of multiple immune and non-immune cell populations in labor-mediator signaling pathways.

Taken together, these findings highlight the complex cell-cell communication network taking place in the murine uterus and how such interactions are modulated by the inflammatory process of preterm labor in both immune and non-immune cell types.

#### Cell-cell communication in the decidua

We next examined the correlations across preterm labor-associated changes in gene expression for each pair of cell types identified in the decidua (Figure S7A). Similar to the uterine tissues, the strongest correlations were observed for non-immune cell types (e.g., stromal, epithelial, fibroblast, smooth muscle, and Endothelial), followed by innate and adaptive immune cells, of which the macrophage and Monocyte clusters were best correlated (Figure S7A). Similarly, the weakest correlations were observed for some epithelial cell types and Neutrophils (Figure S7A). Thus, decidual cells display preterm labor-associated changes in gene expression with varying magnitude of sharing among cell types, which resemble those observed in the uterine tissues.

The inferred cell-cell signaling pathways that were enriched or diminished in the decidua with preterm labor are shown in Figure S7B and Table S3. From among these, the top pathways are displayed with their participating sender and receiver cell types in Figure 5A&B. Similar to the uterine tissues, key cell-cell communication pathways were primarily related to immune functions such as cytokine and chemokine signaling (Figure 5A&B). Unique among the three compared tissues, the IL-17 pathway was most prominent in the decidua (Figure 5A&B), which was consistent with a previous report of IL-17 signaling in endothelial cells derived from the human peripartum decidua (Huang et al., 2020) and suggests that decidual T cells participate in the local inflammatory response to intra-amniotic infection. Among other identified signaling pathways, both immune and non-immune cell subsets contributed as senders or receivers, including the NK-cell-2 subset that was not implicated in uterine cell-cell signaling (Figure 5A&B). The decidual Epithelial-5 cell type appeared to be primarily functioning as a receiver of cell-cell signaling in this tissue (Figure 5A&B). We then visualized the preterm labor-driven changes in incoming and outgoing signaling, and observed that subsets such as Monocyte, Macrophage-1, Neutrophil, NK cells, and Dendritic cells showed predominantly incoming interactions (Figure 5C). On the other hand, stromal and fibroblast subsets as well as T cells tended towards increased outgoing signaling, while SMC-1 and Endothelial cells showed an overall reduction in interaction strength (Figure 5C). Interestingly, outgoing interaction strength was greater in decidual T cells compared to uterine T cells (Figure 5C vs. Figure 4C), which further emphasizes a role for T cell-derived signals in the pathophysiology of preterm labor associated with intra-amniotic infection (Gershater et al., 2022). Consistent with enhanced cell-cell signaling in preterm labor, aggregated cellular interaction plots demonstrated an overall net increase in decidual intercellular interactions compared to controls (Figure 5D). Similar to the uterine tissues, enriched signaling pathways such as CCL and Galectin were primarily driven by Macrophage-1, Monocyte, and Dendritic cell in preterm labor, with overall interactions among cell types increasing compared to controls (Figure 5E).

**Figure 5.**
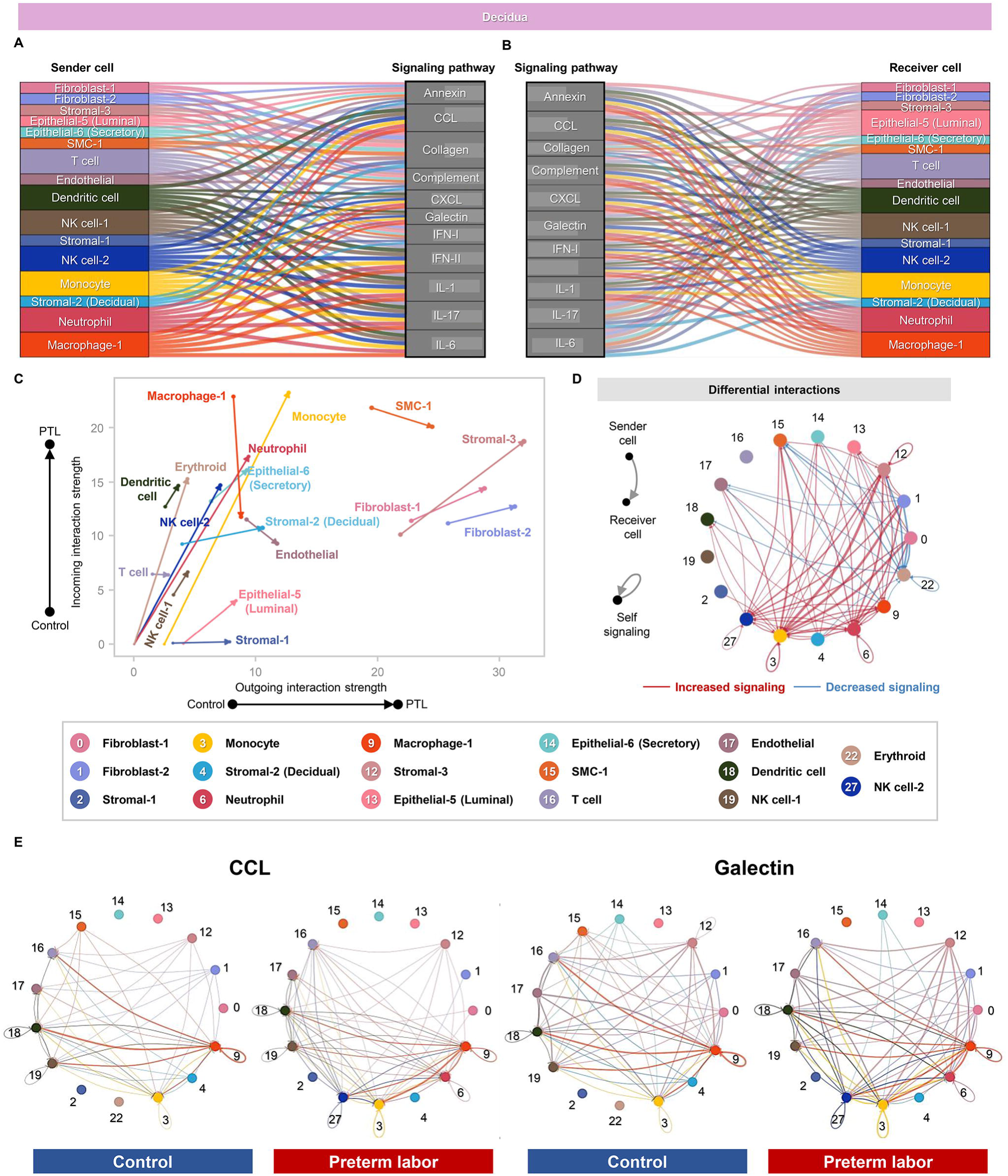
Cellular interactions in the decidua during preterm labor. **(A-B)** Alluvial plots showing the roles of specific cell types as senders or receivers of preterm labor-associated signaling in the decidua based on 11 enriched signaling pathways comparing the overall information flow within networks between preterm labor and control derived from CellChat (full list of pathways in Figure S6B). Connecting lines are color-coded and represent the participation of each cell type as senders or receivers of the indicating signaling pathway. Line thickness is proportional to the strength of interaction. **(C)** Arrow plot showing changes in the outgoing and incoming interaction strength between preterm labor (point of the arrow) and control condition (base of the arrow) for specific cell types in the decidua. **(D)** Circle plots showing the top 25% of aggregated interactions among cell types in the decidua for control and preterm labor groups. Each node represents a cell type and the interaction is shown by color-coded lines. **(E)** Circle plots showing the top 25% increased (red) or decreased (blue) signaling interactions in the decidua for specific pathways in preterm labor compared to controls.

Similar to the changes observed in the uterus, decidual expression of progesterone-related genes was consistently downregulated across cell types with preterm labor (Figure S7C). The patterns of change in prostaglandin-related gene expression were also similar between the decidua and uterus; yet, some difference in the magnitude of change between compartments were observed for multiple genes, potentially indicating a stronger upregulation of preterm labor-associated prostaglandin signaling in the uterus relative to the decidua (Figure S6B).

These data provide insight into the distinct cellular interactions taking place in the decidua during the process of preterm labor, including the involvement of cell types and signaling pathways not observed in other tissues. Yet, the decidua and uterus also share cell type-specific communications that are affected by preterm labor.

#### Cell-cell communication in the cervix

Our investigation of the cell type-specific changes taking place in the cervix with preterm labor had indicated that the Neutrophil and Monocyte subsets were most affected, in tandem with previous studies showing labor-associated infiltration of immune cells (Gonzalez et al., 2011; Osman et al., 2003; Payne et al., 2012; Sakamoto et al., 2005; Timmons et al., 2009), as well as multiple epithelial cell subsets (Figures 2 and 3). Consistently, correlation analysis of these cell types showed that the strongest associations in gene expression changes driven by preterm labor were observed among epithelial cell types (Figure S8A), while Neutrophils and Monocytes showed modest correlation of genes affected by preterm labor (Figure S8A). Inferred cell-cell signaling pathways were noticeably fewer compared to the other tissues and included multiple processes unique to the cervix (Figure S8B and Table S3), which could be attributed to the less diverse cell type composition observed in this tissue. Indeed, as shown by the participating senders and receivers, signaling pathways that were strongly implicated in the uterus and decidua with preterm labor were not as enriched in cervical cell types (Figure 6A&B). On the other hand, cell-cell signaling pathways related to extracellular matrix were strongly represented (Figure 6A&B), which is consistent with the primarily connective tissue composition of the cervix (Granstrom et al., 1989; Krantz and Phillips, 1962; Ludmir and Sehdev, 2000). As expected, given their inferred roles as receivers, most cervical epithelial cell types showed strong incoming interactions with preterm labor, whereas the SMC-1, Fibroblast-2, and Stromal-1 subsets showed a tendency towards increased outgoing interactions (Figure 6B&C). This was further supported by the aggregated cervical cell-cell interactions in the control and preterm labor groups showing increased receipt of signaling by Epithelial-1 and Epithelial-8 as well as SMC-1 and Fibroblast-2 (Figure 6D). Notably, Fibroblast-2 and SMC-1 were top contributors to enriched signaling pathways such as Collagen and Tenascin (Figure 6E). It is possible that the Fibroblast-2 and/or SMC-1 cell clusters include cervical myofibroblasts, given that a previous histological investigation indicated a pregnancy-specific accumulation of such cells, which could be interacting with the extracellular matrix to aid in supporting the mechanical stresses present during labor (Montes et al., 2002). In the last decade, a new paradigm for the role of SMCs in the human cervix has emerged, suggesting a sphincter-like function of the internal os, in which the SMCs express contractility-associated proteins that are responsive to oxytocin signaling (Vink et al., 2016). Together with our current findings, these observations support the involvement of SMC-1 and Fibroblast-2 subsets in preterm labor-associated signaling in the murine cervix, and further emphasize the unique cell-cell signaling pathways taking place in the cervix during preterm labor.

**Figure 6.**
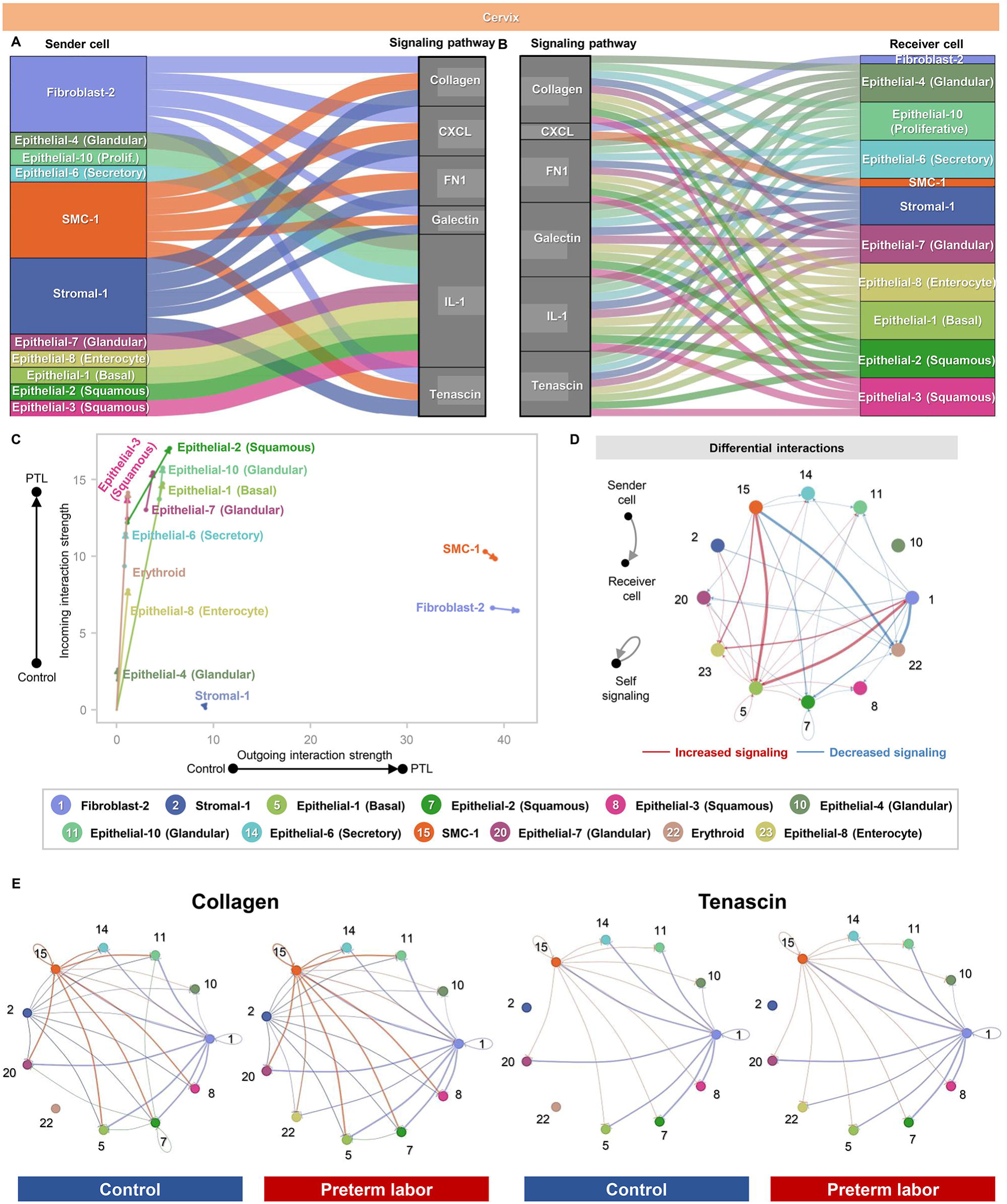
Cellular interactions in the cervix during preterm labor. **(A-B)** Alluvial plots showing the roles of specific cell types as senders or receivers of preterm labor-associated signaling in the cervix based on selected enriched signaling pathways (full list of pathways in Figure S7B). Connecting lines are color-coded and represent the participation of each cell type as senders or receivers of the indicating signaling pathway. Line thickness is proportional to the strength of interaction. **(C)** Arrow plot showing changes in the strength of outgoing and incoming interactions between preterm labor (point of the arrow) and control (base of the arrow) for specific cell types in the cervix. **(D)** Circle plots showing the top 25% of aggregated interactions among cell types in the cervix for control and preterm labor groups. Each node represents a cell type and the interaction is shown by color-coded lines. **(E)** Circle plots showing the top 25% increased (red) or decreased (blue) signaling interactions in the cervix for specific pathways in preterm labor compared to controls.

### Shared cellular signaling pathways in the murine uterus and human myometrium during the processes of preterm and term labor

Last, to examine the shared pathways implicated in the process of parturition in mice and humans, we utilized the differential cellular interactions in the murine uterus with preterm labor together with our previously generated single-cell atlas of the human myometrium with labor at term (Pique-Regi et al., 2022) (Figure 7). We investigated the interaction strength between cell types affected by labor as well as prominent signaling pathways, and contrasted these between the murine and human tissues. Overall, we found that labor-associated cell-cell interactions were primarily driven by SMC, stromal, fibroblast, and innate immune cell types in both the murine uterus and human myometrium, independently of differences in sender/receiver status (Figure 7A&B).

**Figure 7.**
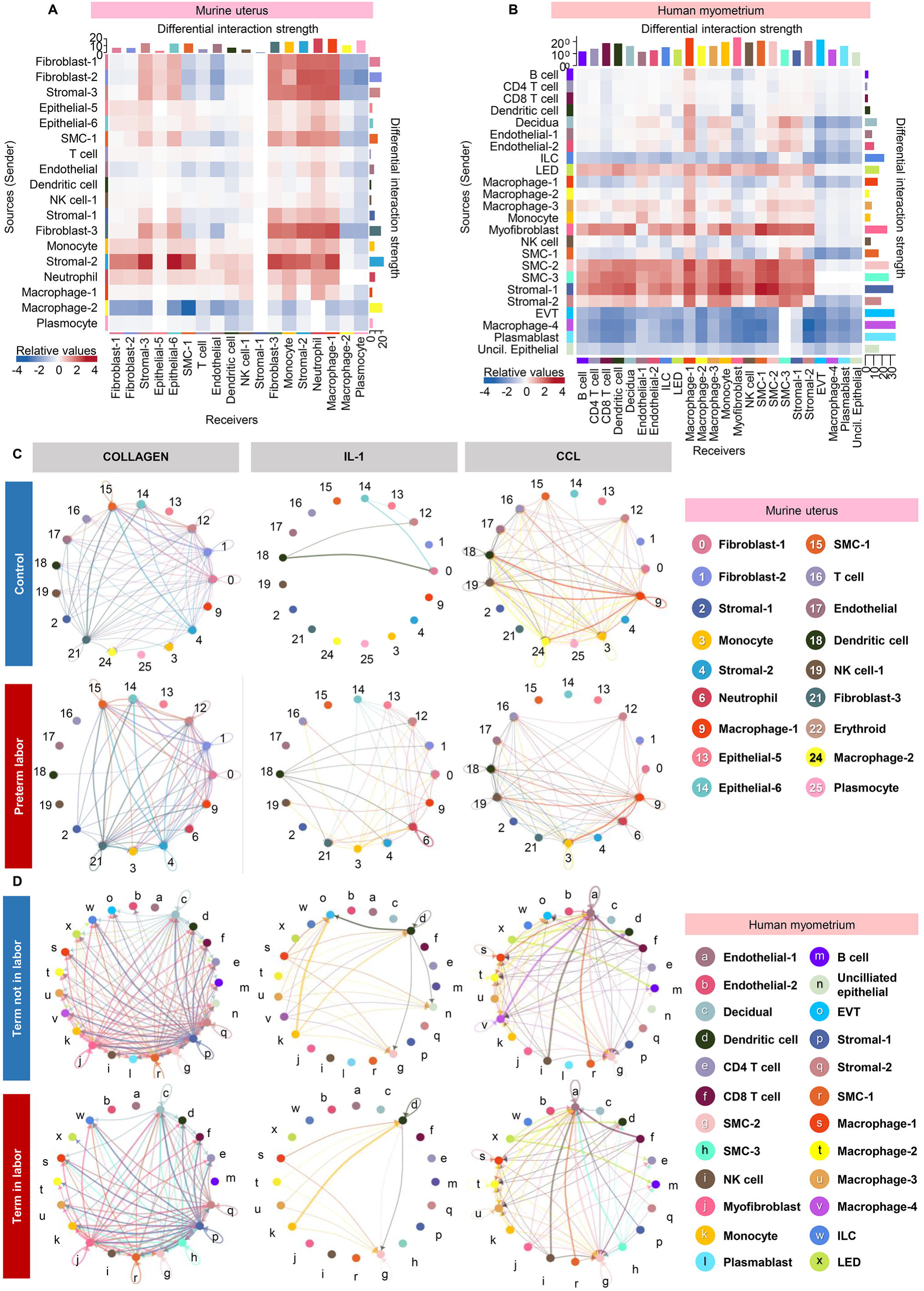
Shared cellular signaling pathways in the murine uterus and human myometrium during the processes of preterm and term labor. **(A)** Heatmap showing the differential interaction strength among cell types in the murine uterus with preterm labor. Red and blue shading indicate increased or decreased signaling, respectively, in preterm labor compared to control. **(B)** Heatmap showing the differential interaction strength among cell types in the human myometrium with term labor. Red and blue shading indicate increased or decreased signaling, respectively, in term labor compared to term without labor. **(C)** Circle plots representing the top 25% murine uterine cell-cell communications inferred for the Collagen, IL-1, CCL pathways for the control and preterm labor groups. **(D)** Circle plots representing the top 25% human myometrial cell-cell communications inferred for the Collagen, IL-1, and CCL pathways for the control and preterm labor groups. Abbreviations used: SMC, smooth muscle cell; NK cell, natural killer cell; EVT, extravillous trophoblast; ILC, innate lymphoid cell; LED, lymphoid endothelial decidual cell.

Specifically, in the murine uterus, non-immune cell types such as Fibroblast-1, -2, and -3, Stromal-2 and -3, and SMC-1 showed the strongest labor-associated increase in outgoing cell-cell signaling, whereas outgoing signaling by Macrophage-2 was greatly diminished (Figure 7A). The top receivers of labor-associated signaling were Fibroblast-3, Stromal-2 and -3, and Epithelial-6 as well as innate immune cell types (Monocyte, Neutrophil, and Macrophage-1) (Figure 7A). Endothelial, SMC-1, Plasmocyte, and Macrophage-2 showed diminished incoming signaling with preterm labor (Figure 7A).

In the human myometrium, labor involved increased outgoing signaling almost exclusively by smooth muscle cell and stromal subsets, with Myofibroblast and LED cells also contributing to this process (Figure 7B). Receivers of such outgoing interactions included macrophage subsets and Monocytes, which is consistent with previous investigations indicating that myometrial cell contraction during labor is promoted by crosstalk with macrophages in a co-culture model (Wendremaire et al., 2020). Multiple myometrial cell subsets displayed substantially reduced outgoing signaling with labor, such as EVT, Macrophage-4, Plasmablast, Unciliated Epithelial, and ILC (Figure 7B). For incoming signaling, the majority of human myometrial cell types tended to have increased interaction only with those cells that displayed greater outgoing signals, such as smooth muscle and stromal cells (Figure 7B). Some exceptions seemed to be the Macrophage-1, SMC-3, and Stromal-1 subsets, whose incoming signaling from the majority of other cell types was strengthened (Figure 7B).

We then examined the top 25% of aggregated cell-cell interactions in the human myometrium at term without labor and at term with labor (Figure S8A). Consistent with the correlation analysis above (Figure 7B), a clear shift in the cell types contributing to myometrial cell-cell signaling was observed between groups, with the EVT, Macrophage-4, Plasmablast, and Unciliated epithelial clusters showing greatly diminished interactions in labor, and smooth muscle and stromal cell subsets acquiring increased interactions (Figure S9A). This shift in interaction with labor was also reflected by the combined differential interaction analysis shown in Figure S8B. When examining the specific cell-cell signaling pathways that were impacted by labor, terms such as Collagen, IL-1, CCL, Complement, and CXCL were found to be shared between the murine uterus and human myometrium, indicating shared labor-associated cellular signaling between species (Figure 7C&D and Figure S9C-G); yet, the closer inspection of these shared signaling pathways revealed differences in the cell types contributing to each (Figure 7C&D and Figure S9C-G). The relevance of immune pathways such as chemokine signaling in the process of labor is supported by previous reports demonstrating that the use of a chemokine inhibitor on human myometrial cells decreased contraction and gap junction formation, thereby disrupting intercellular communication (Boros-Rausch et al., 2021; Coleman et al., 2020). Taken together, these data indicate that labor-associated cell-cell interactions heavily involve SMC, stromal, fibroblast, and innate immune cell types in the murine uterus and the human myometrium, providing evidence of labor-specific signaling processes between immune and non-immune cells that are shared between species. Yet, this interpretation should be taken with caution, given that we compared the physiologic process of labor in the human myometrium with the pathologic process of labor induced by bacteria in the murine uterus.

## DISCUSSION

The current study provides a single-cell atlas of the murine uterus, decidua, and cervix that highlights the cell type composition, transcriptional profiles, and cell-cell signaling taking place in these tissues in normal late gestation or in the context of infection-induced preterm labor and birth. scRNA-seq allowed for the deep characterization of immune and non-immune cells that demonstrates the underappreciated heterogeneity of cell types conventionally considered to be uniform in function, such as uterine smooth muscle cells and cervical epithelial cells. Our data can thus serve as a reference for future studies seeking to target specific subsets of these cells, which may have differing roles in pregnancy and labor as indicated by their distinct transcriptional profiles. Shared modulation of gene expression was noted between uterine and decidual cell types and was reflected by the similar enrichment of labor-associated signaling pathways, which is consistent with the spatial proximity of these tissues. Yet, the comparison of individual cell types across tissues indicated that the most represented biological processes can vary according to location, and therefore the tissue of origin should be taken into consideration when inferring cellular function. Importantly, here we provide scRNA-seq characterization of the understudied cervical tissues, demonstrating a plethora of epithelial subsets with different potential functions as well as smooth muscle and fibroblast cell types that indicate an unexpected level of heterogeneity in the cervix. Recent evidence has suggested a sphincter-like function of SMCs in the internal os of the human cervix (Vink et al., 2016), and our current findings support this concept. Inferred cell-cell communications provided evidence of substantial crosstalk among uterine, decidual, and cervical cell types during the process of preterm labor, highlighting key signaling pathways that could potentially be targeted in future translational studies aimed at preventing spontaneous preterm labor. In addition, such analysis demonstrated cell types with elevated or diminished interactions driven by inflammation, which can serve to identify those cell types that are most and least involved in such signaling. To demonstrate the application of our single-cell dataset, we have leveraged prior single-cell analysis of the human myometrium in term labor to evaluate cellular interactions compared to our murine model of preterm labor. This analysis provided useful insight into shared signaling pathways associated with the inflammatory process of labor, providing a practical demonstration of how our scRNA-seq dataset can be leveraged for *in silico* discovery of specific cell types, pathways, or genes that can be subsequently targeted *in vitro* and/or *in vivo*.

The current study has some limitations. First, it is important to consider that spontaneous preterm labor is a syndrome, for which intra-amniotic infection represents only one known etiology (Romero et al., 2014a). Here, we focused on preterm labor triggered by intra-amniotic infection of *E. coli*, a Gram positive bacteria, using an established animal model that resembles the clinical condition (Gomez-Lopez et al., 2018a). Other known or proposed etiologies for spontaneous preterm labor may have subtle differences in the involved cell types and associated signaling pathways, and thus further characterization of the cellular atlas in each preterm labor subtype is necessary for their distinction. Second, intra-amniotic infection is often polymicrobial and/or can be induced by a variety of bacterial species (Burnham et al., 2020; Romero et al., 2015c), of which *Ureaplasma* species are the most commonly found in the amniotic cavity (Motomura et al., 2020b; Romero et al., 1989; Sweeney et al., 2016; Viscardi, 2010; Yoneda et al., 2016b; Yoon et al., 1998); therefore, the immune responses triggered by each bacterium or cluster of bacteria may differentially affect cellular responses in the reproductive and gestational tissues. However, the *in vivo* standardization of a polymicrobial infection model and the isolation of clinically-relevant *Ureaplasma* species are challenging, and thus here we utilized intra-amniotic infection with an easily-cultured bacterium, *E. coli*, to induce preterm birth in mice. Future investigations may utilize other bacteria detected in the amniotic cavity of women diagnosed with intra-amniotic infection (DiGiulio et al., 2010; Romero et al., 2015c). In addition, it should be noted that single-cell RNA-seq is a discovery-based approach that we have utilized to generate an atlas of the murine reproductive tissues, and thus careful interpretation is required when extrapolating specific findings to the human context. Last, it is worth mentioning that RNA techniques using single-cell suspensions lose information about the spatial relationships among cell types in target tissues; therefore, such data may be complemented by using spatial transcriptomics and/or proteomics. Nonetheless, our data can serve as a resource for targeted studies that can validate such findings using human samples.

## Supporting information

Key Resources Table

## ACKNOWLEDGEMENTS

The authors would like to acknowledge Yesong Liu for his assistance in performing some of the immunofluorescence imaging.

This research was supported, in part, by the Perinatology Research Branch, Division of Obstetrics and Maternal-Fetal Medicine, Division of Intramural Research, *Eunice Kennedy Shriver* National Institute of Child Health and Human Development, National Institutes of Health, U.S. Department of Health and Human Services (NICHD/NIH/DHHS); and, in part, with Federal funds from NICHD/NIH/DHHS under Contract No. HHSN275201300006C. Dr. Romero has contributed to this work as part of his official duties as an employee of the United States Federal Government. This research was also supported by the Wayne State University Perinatal Initiative in Maternal, Perinatal and Child Health. The funders had no role in the study design, data collection and analysis, decision to publish, or preparation of the manuscript. Figures include art created with BioRender.com

## AUTHOR CONTRIBUTIONS

Conceptualization, R.R., R.P-R., and N.G-L.; Methodology, V.G-F., J.G., Y.X., and N.G-L.; Validation, A.P., R.P-R., and A.L.T.; Formal Analysis, V.G-F., A.P., R.P-R., and N.G-L.; Investigation, V.G-F., J.G., E.P., B.P., D.M., Y.X., L.T., Z.L., and A.L.T.; Resources, R.R., A.L.T., R.P-R., and N.G-L.; Data Curation, V.G-F., A.P., and R.P-R.; Writing - Original Draft, V.G-F., R.R., A.P., J.G., B.P., D.M.; Writing - Review & Editing, V.G-F., R.R., A.P., J.G., E.P., B.P., D.M., Y.X., L.T., Z.L., A.L.T, R.P-R., and N.G-L.; Visualization, V.G-F., A.P., J.G., E.P.; Supervision, R.R., R.P-R., and N.G-L.; Project Administration, N.G-L.; Funding Acquisition, R.R., and N.G-L.

## STAR METHODS

### KEY RESOURCES TABLE

(Submitted separately)

### RESOURCE AVAILABILITY

#### Lead contact

Further information and requests for resources and reagents should be directed to and will be fulfilled by the lead contact, Nardhy Gomez-Lopez.

#### Material Availability

This study did not generate new unique reagents.

#### Data and Code Availability

The scRNA-seq data reported in this study has been submitted to the Gene Expression Omnibus repository (GEO accession number GSE200289). All software and R packages used herein are detailed in materials and methods section. Scripts detailing the single-cell analysis are available at GitHub (https://github.com/piquelab/murine_scRNA)

### EXPERIMENTAL MODEL DETAILS

#### Mice

Mice were purchased from The Jackson Laboratory (Bar Harbor, ME, USA) and bred in the animal care facility at the C.S. Mott Center for Human Growth and Development, Wayne State University (Detroit, MI, USA). Mice were under a circadian cycle of light:dark = 12:12 h. Eight-to twelve-week-old C57BL/6 (RRID:IMSR_JAX:000664) female mice were bred with BALB/cByJ male mice (RRID:IMSR_JAX:001026), and females were examined daily between 0800 and 0900 to check for the presence of a vaginal plug, which was considered as 0.5 days *post coitum* (dpc). Upon observation of a vaginal plug, females were removed from mating cages and housed separately, and their weights were monitored daily. At 12.5 dpc, a weight gain of ≥2g was considered confirmation of pregnancy. Mice were randomized to receive intra-amniotic injection of *E. coli* or vehicle control, and investigators were not blinded to control or treatment assignment. No experimental mice were excluded from analysis. All procedures and experiments were approved by the Institutional Animal Care and Use Committee (IACUC) at Wayne State University under Protocol nos. 18-03-0584 and 21-04-3506.

### METHOD DETAILS

#### Preparation of E. coli for intra-amniotic injection

*Escherichia coli* was purchased from the American Type Culture Collection (ATCC, Manassas, VA, USA) (ATCC 12014) and was grown in Luria-Bertani (LB) broth (cat. no. L8050, Teknova, Hollister, CA, USA) at 37°C. From an overnight culture, a sub-culture was placed with fresh LB broth and grown to the logarithmic phase (OD_600_ 0.9 - 1). Additional dilution was performed using sterile 1X phosphate-buffered saline (PBS, Thermo Fisher Scientific/Gibco, Grand Island, NY, USA) to reach a working concentration of 10 CFU/25µL.

#### Measurement of cervical length by ultrasound

Dams were anesthetized on 16.5 dpc by inhalation of 2% isoflurane (Aerrane; Baxter Healthcare Corporation, Deerfield, IL, USA) and 1 to 2 liters/min of oxygen in an induction chamber. Anesthesia was maintained with a mixture of 1.5 to 2% isoflurane and 1.5 to 2 liters/min of oxygen. Mice were positioned on a heating pad and stabilized with adhesive tape. Fur removal from the abdomen was performed using Nair cream (Church & Dwight Co., Inc., Ewing, NJ, USA). Sterile forceps were utilized to expose the vulva and 200 μL of Sterile Aquasonic® 100 ultrasound transmission gel (Parker laboratories, Fairfield, NJ, USA) was used to fill the vagina to create contrast and allow for clear visualization of the external limit of the uterine cervix (i.e., the external os). A trans-abdominal ultrasound approach was utilized to evaluate the cervix. The transducer was slowly moved toward the lower part of the abdomen and the cervix was positioned in a longitudinal view. The cervical length was measured from the internal to the external os at least three times per mouse, and its average was utilized as the final value for cervical length. This procedure was performed prior to ultrasound-guided injection with either *E. coli* or PBS and repeated 24 h later (on 17.5 dpc) (i.e., prior to tissue collection). The change in cervical length was determined as a percentage by considering the cervical length on 16.5 dpc as 100% and then calculating the percentage of the cervical length on 17.5 dpc.

#### Intra-amniotic inoculation with E. coli

Dams that underwent cervical measurement were maintained on the heating pad under anesthesia as described above. The ultrasound transducer was slowly moved toward the abdomen to localize the amniotic sacs. The syringe with *E. coli* suspension (10 CFU/25µL) was stabilized by a mechanical holder (VisualSonics). Ultrasound-guided intra-amniotic inoculation with *E. coli* was performed in each amniotic sac using a 30G needle (BD PrecisionGlide needle; Becton Dickinson, Franklin Lakes, NJ, USA). Controls were injected with 25 μL of sterile 1X PBS into each amniotic sac. After the ultrasound injection, the dams were placed under a heat lamp for recovery (defined as when the mouse resumed normal activities such as walking and responding), which typically occurred 10Lmin after removal from anesthesia. After recovery, mice were video monitored to observe pregnancy outcomes.

#### Video Monitoring

Pregnancy outcomes were recorded via video camera (Sony Corporation, Tokyo, Japan) to determine gestational length, and therefore rate of preterm birth. Preterm birth was defined as delivery occurring before 18.5 dpc, based on the earliest delivery of PBS-injected control dams, and its rate was represented by the percentage of females delivering preterm among the total number of mice injected.

#### Tissue collection prior to preterm birth

Dams were euthanized on 17.5 dpc and the reproductive tissues (uterus, decidua, and cervix) were collected. Tissues collected for the preparation of single-cell suspensions were placed in sterile 1X PBS, while tissues for histological analyses were fixed in 10% Neutral Buffered Formalin (Surgipath, Leyca Biosystems, Wetzlar, Germany) and embedded in paraffin. Five-μm-thick sections were cut and mounted on Superfrost® Plus microscope slides (Cat. No. 48311-703, VWR International, LLC. Radnor, PA, USA).

#### Histological characterization of murine reproductive tissues

##### Leukocyte detection using DAB immunohistochemistry

Five-µm-thick tissue sections from mice injected with PBS or *E. coli* were deparaffinized and rehydrated using xylene and a series of decreasing ethanol concentrations, respectively. Immunohistochemistry staining using the Monoclonal Rabbit Anti-Mouse CD45 (AB_2799780; clone D3F8Q, cat. no. 70257S, Cell Signaling Technology, Danvers, MA, USA) was performed using the Leica Bond Max Automatic Staining System in a peroxidase-mediated oxidation of 3,30-diaminobenzidine (DAB) from the Bond™ Polymer Refine Detection Kit (both from Leica Microsystems, Wetzlar, Germany). The negative control used was the Rabbit FLEX Universal Negative Control (cat. no. IR60066-2, Agilent, Santa Clara, CA, USA). Images were scanned using the Brightfield setting of the Vectra Polaris Multispectral Imaging System.

##### Movat’s pentachrome staining

Five-µm-thick tissue sections from mice injected with PBS or *E. coli* were histologically characterized for the presence of collagen, elastin, muscle, and mucin using the Movat Pentachrome Stain Kit (Modified Russell-Movat; ScyTek Laboratories, Inc. Logan, UT, USA), following manufacturer’s instructions with modifications. Briefly, tissue sections were deparaffinized, stained with working Elastic Stain solution for 20 min, and rinsed in running tap water for 1 min followed by rinsing with deionized water. Then, the following reagents from the kit were sequentially applied to the entire tissue section with distilled water rinsing in between each application: 2% Ferric Chloride for 5-8 s, 5% Sodium Thiosulfate Solution for 1 min, Alcian Blue Solution (pH 2.5) for 20 min, Biebrich Scarlet-Acid Fuchsin Solution for 2 min, 5% Phosphotungstic Acid Solution for 7 min, and 1% Acetic Acid Solution for 3 min. Excess Acetic Acid Solution was drained from the slides and Yellow Stain Solution was immediately applied for 20 min. The slides were then rinsed in 100% ethanol followed by rinsing with xylene. Images were scanned using the Brightfield setting of the Vectra Polaris Multispectral Imaging System (Akoya Biosciences, Marlborough, MA, USA).

##### OPAL multiplex immunofluorescence

OPAL multiplex immunofluorescence staining was performed using the OPAL Multiplex 7-color IHC kit (Cat. no. NEL811001KT; Akoya Biosciences), according to the manufacturer’s instructions. Prior to multiplex staining, the order of antibody staining was optimized using single-plex staining paired with tyramide signal amplification (TSA)-conjugated OPAL fluorophores. The optimized detection panel includes antibody-OPAL fluorophore pairs in the following order: Monoclonal Rabbit Anti-Mouse F4/80 (AB_2799771; clone D2S9R; cat. no. 70076S, Cell Signaling Technology) with OPAL 520, Monoclonal Rabbit Anti-Mouse CD3ε (AB_2889902; clone E4T1B; cat. no. 78588S, Cell Signaling Technology) with OPAL 570, Monoclonal Rabbit Anti-Mouse Klrb1c/CD161c (AB_2892989; clone E6Y9G; cat. no. 39197S, Cell Signaling Technology) with OPAL 620, Polyclonal Rabbit Anti-Mouse Ly6C (cat. no. HA500088, HuaBio, Boston, MA, USA) with OPAL 650, and Monoclonal Rabbit Anti-Mouse Ly6G (AB_2909808; clone E6Z1T; cat. no. 87048S, Cell Signaling Technology) with OPAL 690. The Rabbit FLEX Universal Negative Control (Agilent) was used as isotype. Briefly, 5-µm-thick tissue sections from mice injected with PBS or *E. coli* were deparaffinized and rehydrated using xylene and a series of decreasing ethanol concentrations, respectively. The slides were rinsed in deionized water and epitope retrieval was performed by submerging the slides in appropriate antigen retrieval (AR) buffer and boiling in a microwave oven. Non-specific binding was prevented by incubating slides in OPAL antibody diluent/blocking solution prior to incubating with each primary antibody at room temperature. Next, the slides were rinsed in TBST prior to incubation with anti-mouse secondary antibody-horse radish peroxidase (HRP) conjugate followed by the selected TSA-conjugated OPAL fluorophore. Cycles of sequential epitope retrieval, target detection, and signal amplification were repeated using the optimized antibody-OPAL fluorophore pair. Once all targets were detected, the slides were incubated with DAPI (4′,6-diamidino-2-phenylindole) as a nuclear counterstain and mounted using AquaSlip™ Aqueous Permanent Mounting Medium (American MasterTech). Fluorescence image acquisition was performed using the Vectra Polaris Multispectral Imaging System at 20x magnification. Multispectral images were analyzed using the inForm software version 2.4 (Akoya Biosciences).

#### Tissue dissociation of murine uterus and decidua

Immediately following tissue collection, the uterus and decidua were dissociated to prepare single-cell suspensions. The tissues were mechanically dissociated and enzymatically digested by incubating at 37°C using enzymes from the Umbilical Cord Dissociation Kit (Miltenyi Biotec). A second round of mechanical dissociation was performed using the gentleMACS Dissociator (Miltenyi Biotec), and dissociated cells were rinsed with 1X PBS (Thermo Fisher Scientific) prior to filtration using a 100µm cell strainer (Miltenyi Biotec). Filtered cells were pelleted by centrifugation at 300 x g for 5 min, erythrocytes were eliminated using ACK Lysing Buffer (Life Technologies), and the cells were rinsed in 0.04% BSA (Sigma Aldrich) and 0.5 mM EDTA (Sigma Aldrich) diluted in 1X PBS. Finally, the cells were filtered using a 30µm cell strainer (Miltenyi Biotec), and the Dead Cell Removal Kit was used to remove dead cells to obtain a cell viability of ≥80%.

#### Tissue dissociation of the murine cervix

Immediately following the collection of the cervix, the tissue was mechanically dissociated and enzymatically digested using Collagenase A (160 mg/mL) (Sigma Aldrich, St. Louis, MO, USA) and incubated at 37°C. Then, the dissociated cells were pelleted by centrifugation at 16,000 x g for 10 min at 20°C and resuspended with 0.05% trypsin-EDTA (Thermo Fisher Scientific, Waltham, MA) prior to a second round of mincing and incubation in 0.05% trypsin-EDTA at 37°C. The enzymatic reaction was stopped by the addition of FBS (Fetal Bovine Serum, Thermo Fisher). Cells were then filtered using a 70µm cell strainer (Miltenyi Biotec, San Diego, CA, USA) and pelleted by centrifugation at 300 x g for 10 min. Erythrocytes were removed using ACK Lysing Buffer (Life Technologies, Grand Island, NY, USA). Finally, the cells were resuspended in 0.04% Bovine Serum Albumin (BSA) (Sigma Aldrich) diluted in 1X PBS (Thermo Fisher Scientific, NY, USA) and filtered through a 30µm cell strainer (Miltenyi Biotec). The cell concentration and viability were determined using an automatic cell counter (Cellometer Auto 2000, Nexcelom Bioscience, Lawrence, MA, USA) and the Dead Cell Removal Kit (Miltenyi Biotec) was used to remove dead cells to obtain a cell viability of ≥80%.

#### Generation of gel beads-in-emulsion (GEMs) and library preparation

Generation of gel beads-in-emulsion (GEMs) and preparation of library constructs was performed on viable single-cell suspensions using the 10x Genomics Chromium Single Cell 3’ Gene Expression Version 3.1 Kit (10x Genomics, Pleasanton, CA, USA), according to the manufacturer’s instructions. Briefly, viable single cells were encapsulated in partitioning oil together with a single Gel Bead with barcoded oligonucleotides within the Chromium Controller. Reverse transcription of mRNA into complementary (c)DNA was performed using the Veriti 96-well Thermal Cycler (Thermo Fisher Scientific, Wilmington, DE, USA). Dynabeads MyOne SILANE (10x Genomics) and the SPRIselect Reagent (Beckman Coulter, Indianapolis, IN, USA) were used to purify resulting cDNA, which was optimized by enzymatic fragmentation, end-repair, and A-tailing. Next, adaptors and sample index were incorporated by ligation. The sample index PCR product was then amplified using the Veriti 96-well Thermal Cycler and double-sided size selection was performed using the SPRIselect Reagent. Following the formation of cDNA and final library construct, the Agilent Bioanalyzer High Sensitivity DNA Chip (Agilent Technologies, Wilmington, DE, USA) was used determine sample quality and concentration.

#### Sequencing

Prior to sequencing of post-library constructs, samples were quantified using the Kapa DNA Quantification Kit for Illumina platforms (Kapa Biosystems, Wilmington, MA, USA), following the manufacturer’s instructions. The sequencing of 10x scRNA-seq libraries was performed on the Illumina NextSeq 500 at the Genomics Services Center (GSC) of the Center for Molecular Medicine and Genetics (Wayne State University School of Medicine, Detroit, MI, USA). The Illumina 75 Cycle Sequencing Kit (Illumina, San Diego, CA, USA) was used with 58 cycles for R2, 26 for R1, and 8 for I1.

### QUANTIFICATION AND STATISTICAL ANALYSIS

#### scRNA-seq data normalization and pre-processing

Sequencing data were processed using Cell Ranger version 4.0.0 (10x Genomics). The “cellranger counts” was also used to align the scRNA-seq reads by using the STAR aligner(Dobin et al., 2013) to produce the bam files necessary for demultiplexing the individual of origin based on genotype information using demuxlet(Kang et al., 2018) and a custom vcf file. The genotype data were downloaded from ftp://ftp-mouse.sanger.ac.uk/current_snps/mgp.v5.merged.snps_all.dbSNP142.vcf.gz, the strains C57BL_6NJ and BALB_cJ were extracted, and a new synthetic vcf file was generated consisting of all the genetic variants where these two strains diverge, and containing a maternal genotype column identical to the C57BL_6NJ strain and a fetal genotype column with a “0/1” heterozygote genotype. Ambient RNA contamination and doublets were removed using SoupX version 1.5.2(Young and Behjati, 2020) and DoubletFinder 2.0.3(McGinnis et al., 2019). Additionally, any cell with < 200 genes or > 20,000 genes detected, or that had > 10% mitochondrial reads, was excluded (Table S4). All count data matrices were then normalized and combined using the Seurat package in R (Seurat version 4.0.3)(Hafemeister and Satija, 2019; Stuart et al., 2019). The first 100 principal components were obtained, and the different libraries were integrated and harmonized using the Harmony package in R version 1.0.0(Korsunsky et al., 2019). The top 30 harmony components were then processed to embed and visualize the cells in a two-dimensional map via the Uniform Manifold Approximation and Projection for Dimension Reduction (UMAP) algorithm(Becht et al., 2018; McInnes et al., 2018). A resolution of 0.5 was used to cluster the single cells.

#### Annotation of cell types

The SingleR(Aran et al., 2019) package in R version 1.6.1 was used to annotate cell types based on their similarities to reference datasets with known labels(Buechler et al., 2021; Tabula Muris et al., 2018). SingleR annotates single cells from query data by computing the Spearman’s correlation coefficient between the single-cell gene expression data and samples from the reference dataset. The correlation is measured only based on the variable genes in the reference dataset. The multiple correlation coefficients per cell type are combined according to the cell type labels of the reference dataset to assign a score per cell type. Additionally, we confirmed the cell type identities by identifying the top DEGs (see below) and the gene-cell type mapping data provided by the Mouse Cell Atlas and single-cell MCA (scMCA) package(Han et al., 2018) in R version 0.2.0. Using different annotations obtained from the reference mapping workflows, the final cell type labels were assigned based on a majority vote. If multiple clusters were assigned to the same consensus cell type, we added a sub-index to that cell type for each different original Seurat cluster: Clusters 0, 1, and 21 were annotated as Fibroblast-1, Fibroblast-2, and Fibroblast-3; clusters 2, 4, and 12 were annotated as Stromal-1, Stromal-2 (Decidua), and Stromal-3; clusters 5, 7, 8, 10, 11, 13, 14, 20, 23, and 28 were annotated as Epithelial-1 (basal), Epithelial-2 (squamous), Epithelial-3 (squamous), Epithelial-4 (glandular), Epithelial-10 (proliferative), Epithelial-5 (luminal), Epithelial-6 (secretory), Epithelial-7 (glandular), Epithelial-8 (Enterocyte), and Epithelial-9 (Secretory); clusters 9 and 24 were annotated as Macrophage-1 and Macrophage-2 (progenitor); clusters 15 and 26 were annotated as SMC-1 and SMC-2; and clusters 19 and 27 were annotated as NK-cell-1 and NK-cell-2. All remaining clusters were assigned a unique cell type identifier (Table S5).

#### Differential gene expression for cell type analysis

For this analysis, the differential expression of selected marker genes for each cell type/cluster was identified using the Wilcoxon Rank Sum test and the FindAllMarkers function from Seurat (Table S5). For this analysis, we first compared each cluster to all cell types. We further used the top cell markers [ranked based on log_2_(Fold change) and requiring q < 0.1] assigned to each sub-cluster to annotate the clusters using the Mouse Cell Atlas and scMCA package(Han et al., 2018).

#### Differential gene expression in preterm labor

The identification of preterm labor-associated DEGs between study groups was performed using the DESeq2 R package version 1.32.0(Love et al., 2014). A term for each library was added to the DESeq2 model to correct for technical batch effects (library identifier). For each cell type/replicate combination, we only used combinations with more than 20 cells; otherwise, it was treated as non-observed. Cell types found in < 3 combinations per study group were dropped from the differential gene expression analysis (Table S2 contains all genes determined as differentially expressed). Note that these thresholds imply that clusters with < 120 cells are not analyzed to ensure robust gene expression estimation. Quantile-quantile plots were used to show that p-values are well calibrated under the null hypothesis of no effect of preterm labor, and also to show which tissues and cell types are more enriched for preterm labor-associated gene expression changes (Figure 2H-J). Multiple comparison correction was performed by controlling for false discovery rate using Benjamini-Hochberg’s method and genes with q < 0.1 were reported in Figure 2E-G and Table S2. Statistical difference between the fraction of genes that were upregulated versus downregulated by preterm labor in each cell type was assessed with a binomial test and corrected for multiple comparisons using Benjamini-Hochberg’s method. To compare the effects of preterm labor on gene expression across different tissues and cell types, we performed Spearman’s correlation between the log_2_FC obtained in each DESeq2 analysis performed using genes that had been detected as differentially expressed in at least one cell-type/tissue, q < 0.1. These correlations were visualized as a heatmap in Figure S5A, S6A, and S7A and in boxplots for relevant tissue and cell-type combinations in Figure 3B-D.

#### Gene ontology and pathway enrichment analysis of genes affected by preterm labor

The clusterProfiler in R version 4.0.4(Yu et al., 2012) was used to perform the Over-Representation Analysis (ORA) separately for each list of genes obtained as differentially expressed for each cell type based on the Gene Ontology (GO), Kyoto Encyclopedia of Gene and Genomes (KEGG), and Reactome databases. The functions “enrichPathway”, “enrichKEGG”, and “enrichGO”, from “clusterProfiler” were used. In ORA analyses, the universe of genes for each cell type was the subset that was expressed at a level sufficient to be tested in differential gene expression analysis. When results are combined across cell types, any genes tested (with a calculated p-value) in any of the cell types are used for the universe. Only ORA results that were significant after correction were reported with q < 0.05 being considered statistically significant.

#### Cell-cell communication analysis

CellChat (Jin et al., 2021) was used to infer the cell-cell communications using the single-cell gene expression data from preterm labor and control conditions and a database of prior knowledge of the interactions between signaling ligands, receptors, and their cofactors. The top 25% of significant cell-cell communications (p < 0.05) across different pathways were shown for the two conditions of preterm labor and control. Next, the aggregated cell-cell communication between different cell groups was calculated for the two study groups, and the interaction strength was compared among different cell types from the two study groups. The differential interaction strength was represented with circle plots with red (or blue) edges showing the increased (or decreased) signaling in preterm labor compared to controls. Additionally, the detailed differential interaction strengths were shown using heatmap representations. Major signaling sender and receiver cells were displayed using scatter plots where the changes in signaling strength from control to preterm labor were represented by arrows. The R packages CellChat version 1.1.2, ggalluvial version 0.12.3, and ggplot2 version 3.3.5 were used to visualize cell-cell communication analyses. The major sending and receiving signaling roles based on context-specific pathways across different cell groups were identified using a cut-off of 0.5 when visualizing the connection. The overall information flow [sum of the significant communication probability (p < 0.05) in the inferred cell-cell network] for each signaling network was represented using a bar plot. The comparison between the overall information flow from the two study groups (preterm labor and control) was performed using the paired Wilcoxon test with the function “rankNet” from CellChat.

#### Comparison between cell-cell communication in human and murine uterine tissues

We inferred cell-cell communications using the human myometrial single-cell gene expression data from term in labor (TIL) and term without labor (TNL) study groups(Pique-Regi et al., 2022), and compared the inferred interactions between mouse (uterus) and human (myometrium) across the top common signaling pathways with highest numbers of DEGs.

#### Statistical analysis

Observational mouse data were analyzed by using SPSS v19.0 and GraphPad Prism version 8. For comparing the rates of preterm birth, the Fisher’s exact test was used. For gestational length and cervical shortening, the statistical significance of group comparisons was assessed using the Mann-Whitney U-test or paired t-test, respectively.

## SUPLEMENTAL INFORMATION

### Supplementary Figure Legends

**Figure S1. Preterm labor induced by intra-amniotic infection alters the cellular composition of the murine reproductive tissues, related to Figure 1**. **(A)** UMAP plots showing the cell types present in the uterus, decidua, and cervix of control mice and **(B)** mice with preterm labor. **(C-E)** UMAP plots showing the distribution of cells according to fetal (purple) or maternal (grey) origin in the uterus, decidua, and cervix. Abbreviations used: SMC, smooth muscle cell; NK cell, natural killer cell.

**Figure S2. Leukocyte infiltration in the uterus, decidua, and cervix, related to Figure 2**. **(A-C)** Representative Movat pentachrome staining images of the uterus, decidua, and cervix from control (top row) and preterm labor (bottom row) mice. Red staining indicates muscle/fibrin, dark purple staining indicates elastic fibers, blue staining indicates mucin, and yellow indicates collagen/reticular fibers. Nuclei appear as dark blue/black. Whole-slide images taken at (A) control: 3.7X, preterm labor 4X; (B) control: 2.9X, preterm labor: 4.9X; (C) control: 3.2 X, preterm labor: 3.1X (scale bars = 300μm). Zoomed images were all taken at 20X magnification (scale bars = 50μm). **(D-F)** Representative images showing 3, 3’diaminobenzidine (DAB) immunohistochemistry to detect the pan-leukocyte marker CD45 in the uterus, decidua, and cervix of control mice (top row) and preterm labor (bottom row) mice (n = 3 per group). Whole-slide brightfield images taken at (E) control: 3.5X, preterm labor: 3.3X; (F) control: 2.9X, preterm labor 3.9X; (G) control: 3.4X, preterm labor: 3.2X (scale bars = 300μm). Zoomed images were taken at 20X magnification (scale bars = 50μm). **(G-I)** Representative merged image showing the co-localized immunofluorescence detection of neutrophils (Ly6G+ cells, pink), monocytes (Ly6C+ cells, cyan), macrophages (F4/80+ cells, red), T cells (CD3+ cells, yellow) and NK cells (CD161c+ cells, green) in the uterus, decidua, and cervix of control and preterm labor mice only (n = 3). Nuclear staining is shown in blue (4’,6-diamidino-2-phenylindole; DAPI). Images were taken at 20X magnification. Scale bar = 100μm.

**Figure S3. Shared and unique biological processes in specific cell types impacted by labor across tissues, related to Figure 3**. **(A)** Box plots showing the Spearman’s correlation of the preterm labor-associated log_2_(Fold change, FC) among cell types within each tissue. **(B)** Box plots showing the Spearman’s correlations between tissue pairs for the preterm labor-associated logFC in cell types present in the two tissues. **(C)** ClusterProfiler dot plot showing preterm labor-associated Kyoto Encyclopedia of Genes and Genomes (KEGG) pathways enriched in specific cell 588 types in the uterus, decidua, and cervix. The size and color of each dot represents enrichment score and significance level, respectively. Significant KEGG pathways (q < 0.05) were identified based on over-representation analysis using one-sided Fisher’s exact tests. Abbreviations used: SMC, smooth muscle cell; NK cell, natural killer cell.

**Figure S4. The uterus and decidua share enrichment of biological processes in preterm labor, related to Figure 3**. Cluster profiler dot plots showing the Gene Ontology (GO) biological processes enriched with preterm labor in **(A)** Stromal-1 and Stromal-2, **(B)** Stromal-3 and Fibroblast-1, **(C)** Fibroblast-2 and Fibroblast-3, and **(D)** Endothelial cells in the uterus and decidua. The size and color of each dot represent gene ratio and significance level, respectively. A 1-sided Fisher’s exact test was used.

**Figure S5. Preterm labor-associated enrichment of signaling pathways in uterine cell types, related to Figure 4**. **(A)** Heatmap showing correlations among uterine cell types where red and white blocks signify increased and decreased correlation, respectively. Pearson correlation tests was used. **(B)** Bar plots showing the high expression of specific signaling pathways in the uterus of control mice (blue bars/pathway names) or mice with preterm labor (red bars/pathway names). **(C)** Forest plot showing the log_2_(FC, fold change) and 95% confidence intervals of differentially expressed (DEGs) across cell types in murine uterus. DEGs shown are significant with FDR (q < 0.1). Abbreviations used: SMC, smooth muscle cell; NK cell, natural killer cell.

**Figure S6. Preterm labor-induced induced changes in the expression of prostaglandin-associated genes in different cell types across the reproductive tissues, related to Figures 4–6**. Forest plot showing the log_2_(FC, fold change) and 95% confidence intervals of differentially expressed (DEGs) in selected cell types across the murine cervix, decidua and uterus. DEGs shown are significant with FDR (q < 0.01).

**Figure S7. Preterm labor-induced changes in interaction between decidual cell types, related to Figure 5**. **(A)** Heatmap showing correlations among decidual cell types where red and white blocks signify increased and decreased correlation, respectively. Pearson correlation tests was used. **(B)** Bar plots showing the high expression of specific signaling pathways in the decidua of control mice (blue bars/pathway names) or mice with preterm labor (red bars/pathway names). **(C)** Forest plot showing the log_2_(FC, fold change) and 95% confidence intervals of differentially expressed (DEGs) across cell types in murine decidua. DEGs shown are significant with FDR (q < 0.1). Abbreviations used: SMC, smooth muscle cell; NK cell, natural killer cell.

**Figure S8. Preterm labor-induced changes in interaction between cervical cell types, related to Figure 6**. **(A)** Heatmap showing correlations among cervical cell types where red and white blocks signify increased and decreased correlation, respectively. Pearson correlation tests was used. **(B)** Bar plots showing the high expression of specific signaling pathways in the cervix of control mice (blue bars/pathway names) or mice with preterm labor (red bars/pathway names).

**Figure S9. Labor-associated signaling in the human myometrium partially overlaps with preterm labor-associated changes in the murine uterus, related to Figure 7**. **(A)** Circle plots showing the top aggregated interactions among cell types in the myometrium from humans without (left) or with labor at term (right). Each node represents a cell type and the interaction is shown by lines color-coded based on the sender cell. Representation of aggregated interactions with p < 0.05 using cell chat. **(B)** Circle plot showing the increased (red) or decreased (blue) signaling interactions in the human myometrium in labor compared to controls without labor. Representation of top 25% differential interaction strength. **(C)** Venn diagram showing the overlap in upregulated signaling pathways between the murine uterus in preterm labor (left circle, pink) and the human myometrium in term labor (right circle, orange). Shared labor- and inflammation-associated pathways are highlighted in red. (D-E) Circle plots representing the top 25% human myometrial cell-cell communications inferred for the Complement and CXCL pathways for the Term not in labor and Term in labor groups. (F-G) Circle plots representing the top 25% murine uterine cell-cell communications inferred for the Complement and CXCL pathways for the control and preterm labor groups. Abbreviations used: SMC, smooth muscle cell; NK cell, natural killer cell; EVT, extravillous trophoblast; ILC, innate lymphoid cell; LED, lymphoid endothelial decidual cell.

## SUPPLEMENTAL EXCEL TABLE TITLES

**Table S1.** Summary of the numbers of uterine, decidual and cervical cells encapsulated within the 10x Genomics Gel-bead-in-emulsion (GEM), related to Figure 2.

**Table S2.** Preterm labor-associated differentially expressed genes in uterine, decidual and cervical cell types, as well as the total differentially expressed genes across tissue, related to Figures 2 and 3.

**Table S3.** Summary of genes and pathways used in the CellChat intercellular communication analysis for the uterus, decidua and cervix, related to Figures 4, 5, and 6.

**Table S4.** Summary of quality control metrics calculated with the 10x Genomics Cell Ranger pipeline for each library, related to methods section **“*scRNA-seq data normalization and pre-processing*”**.

**Table S5.** List of marker genes for the identification of cell types in the murine uterus, decidua and cervix, related to methods section **“*Annotation of cell types*”**.

