## Supplementary material for "The single-cell atlas of the murine reproductive tissues during preterm labor": Figure S

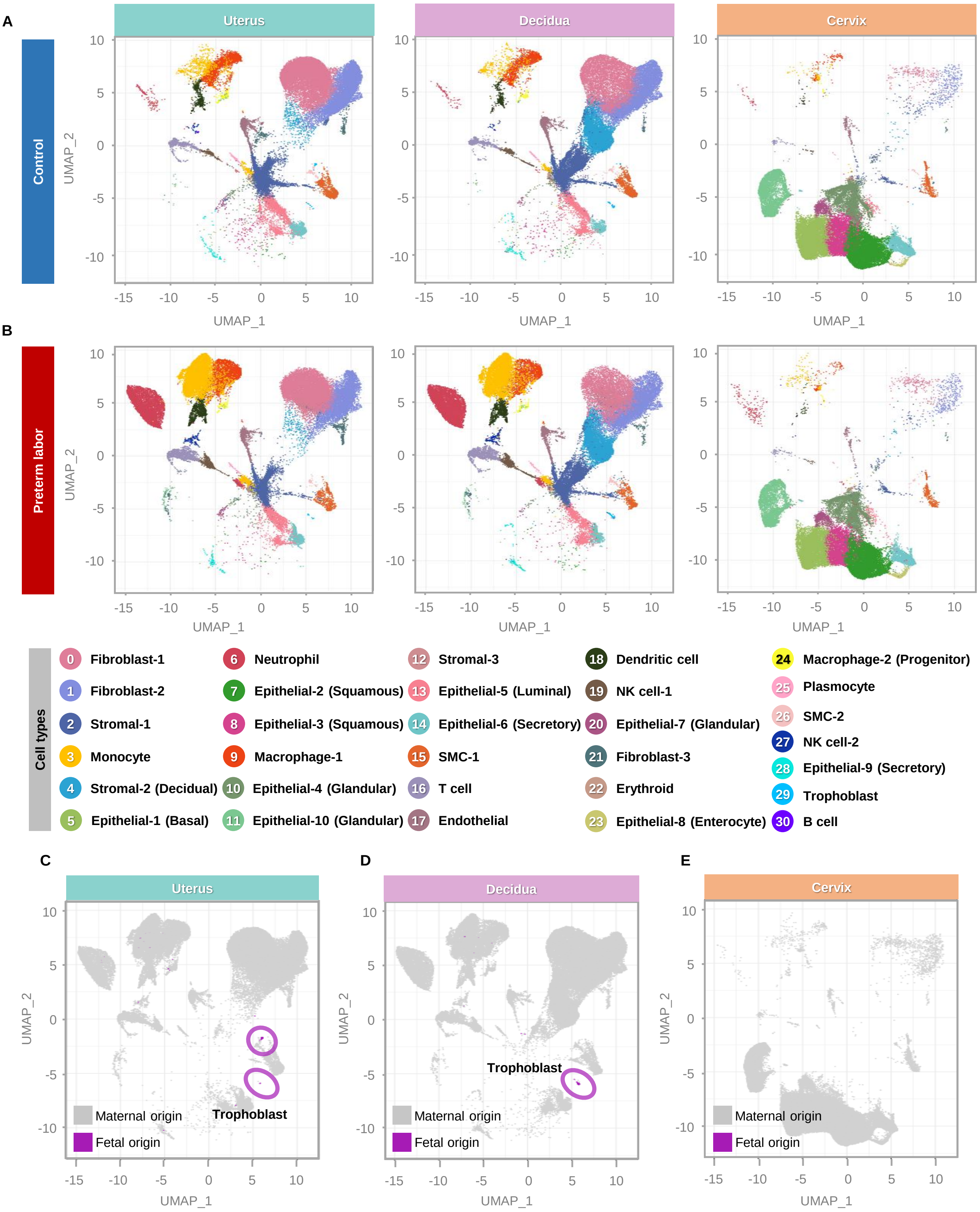


**Figure S1. Preterm labor induced by intra-amniotic infection alters the cellular composition of the murine reproductive tissues, related to Figure 1. (A)** UMAP plots showing the cell types present in the uterus, decidua, and cervix of control mice and **(B)** mice with preterm labor. **(C-E)** UMAP plots showing the distribution of cells according to fetal (purple) or maternal (grey) origin in the uterus, decidua, and cervix. Abbreviations used: SMC, smooth muscle cell; NK cell, natural killer cell.

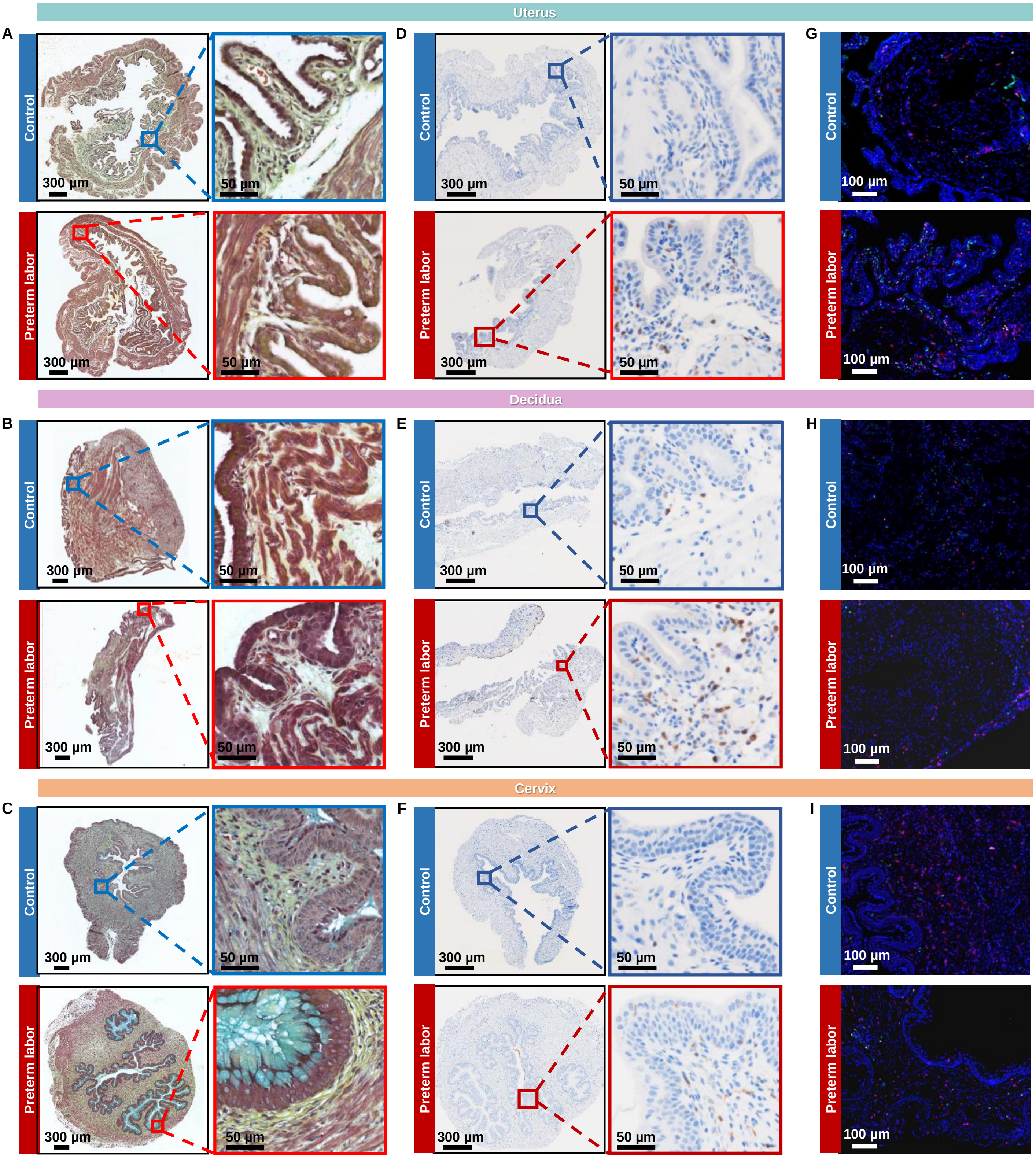


**Figure S2. Leukocyte infiltration in the uterus, decidua, and cervix, related to Figure 2. (A-C)** Representative Movat pentachrome staining images of the uterus, decidua, and cervix from control (top row) and preterm labor (bottom row) mice. Red staining indicates muscle/fibrin, dark purple staining indicates elastic fibers, blue staining indicates mucin, and yellow indicates collagen/reticular fibers. Nuclei appear as dark blue/black. Whole-slide images taken at (A) control: 3.7X, preterm labor 4X; (B) control: 2.9X, preterm labor: 4.9X; (C) control: 3.2 X, preterm labor: 3.1X (scale bars = 300µm). Zoomed images were all taken at 20X magnification (scale bars = 50µm). **(D-F)** Representative images showing 3, 3'diaminobenzidine (DAB) immunohistochemistry to detect the pan-leukocyte marker CD45 in the uterus, decidua, and cervix of control mice (top row) and preterm labor (bottom row) mice (n = 3 per group). Whole-slide brightfield images taken at (E) control: 3.5X, preterm labor: 3.3X; (F) control: 2.9X, preterm labor 3.9X; (G) control: 3.4X, preterm labor: 3.2X (scale bars = 300µm). Zoomed images were taken at 20X magnification (scale bars = 50µm). **(G-I)** Representative merged image showing the co-localized immunofluorescence detection of neutrophils (Ly6G+ cells, pink), monocytes (Ly6C+ cells, cyan), macrophages (F4/80+ cells, red), T cells (CD3+ cells, yellow) and NK cells (CD161c+ cells, green) in the uterus, decidua, and cervix of control and preterm labor mice only (n = 3). Nuclear staining is shown in blue (4',6-diamidino-2-phenylindole; DAPI). Images were taken at 20X magnification. Scale bar = 100µm.

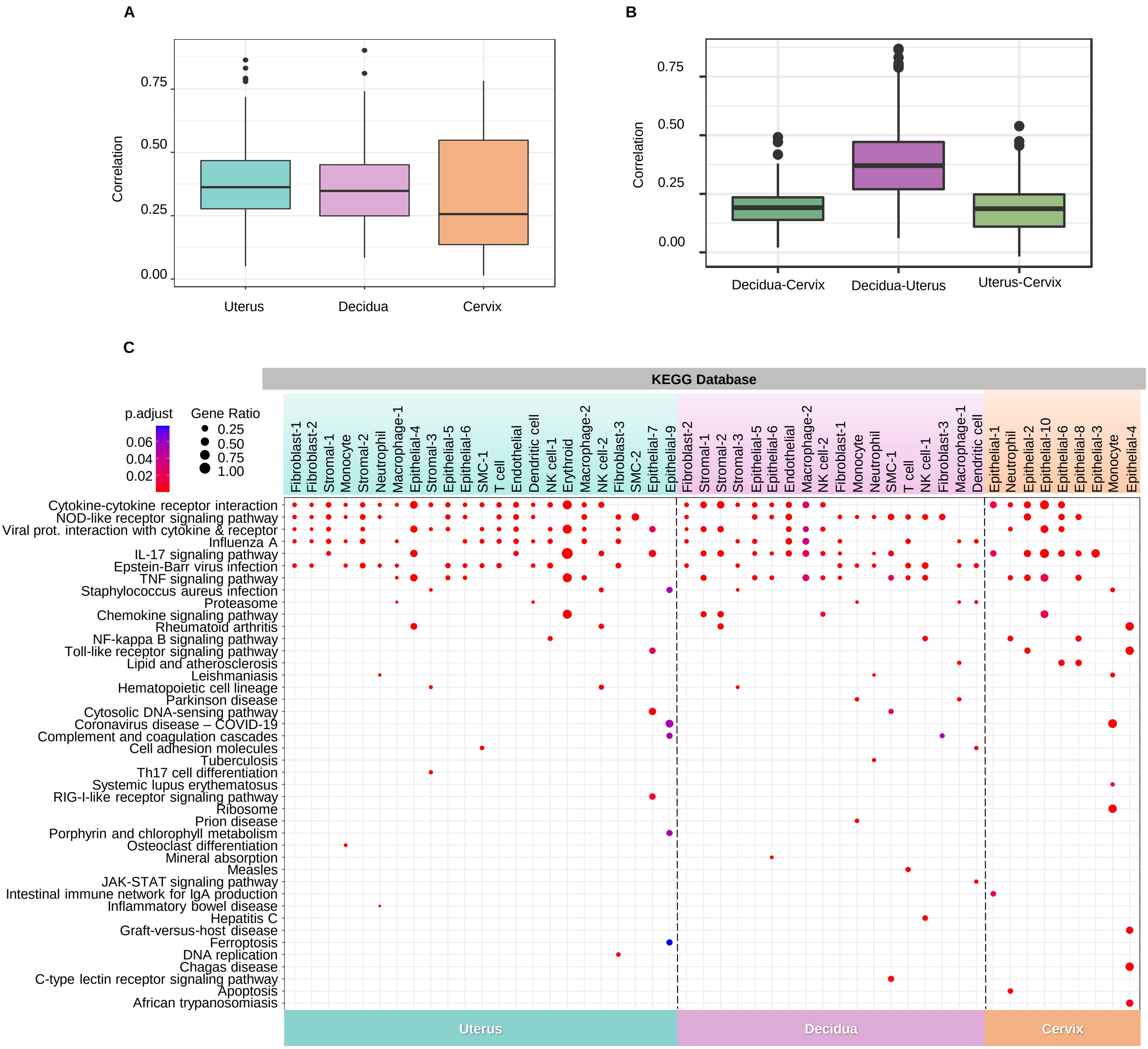


**Figure S3. (A)** Box plots showing the Spearman's correlation of the preterm labor-associated $\log_2$(Fold change, FC) among cell types within each tissue. **(B)** Box plots showing the Spearman's correlations between tissue pairs for the preterm labor-associated logFC in cell types present in the two tissues. **(C)** ClusterProfiler dot plot showing preterm labor-associated Kyoto Encyclopedia of Genes and Genomes (KEGG) pathways enriched in specific cell 588 types in the uterus, decidua, and cervix. The size and color of each dot represents enrichment score and significance level, respectively. Significant KEGG pathways ($q < 0.05$) were identified based on over-representation analysis using one-sided Fisher's exact tests. Abbreviations used: SMC, smooth muscle cell; NK cell, natural killer cell.

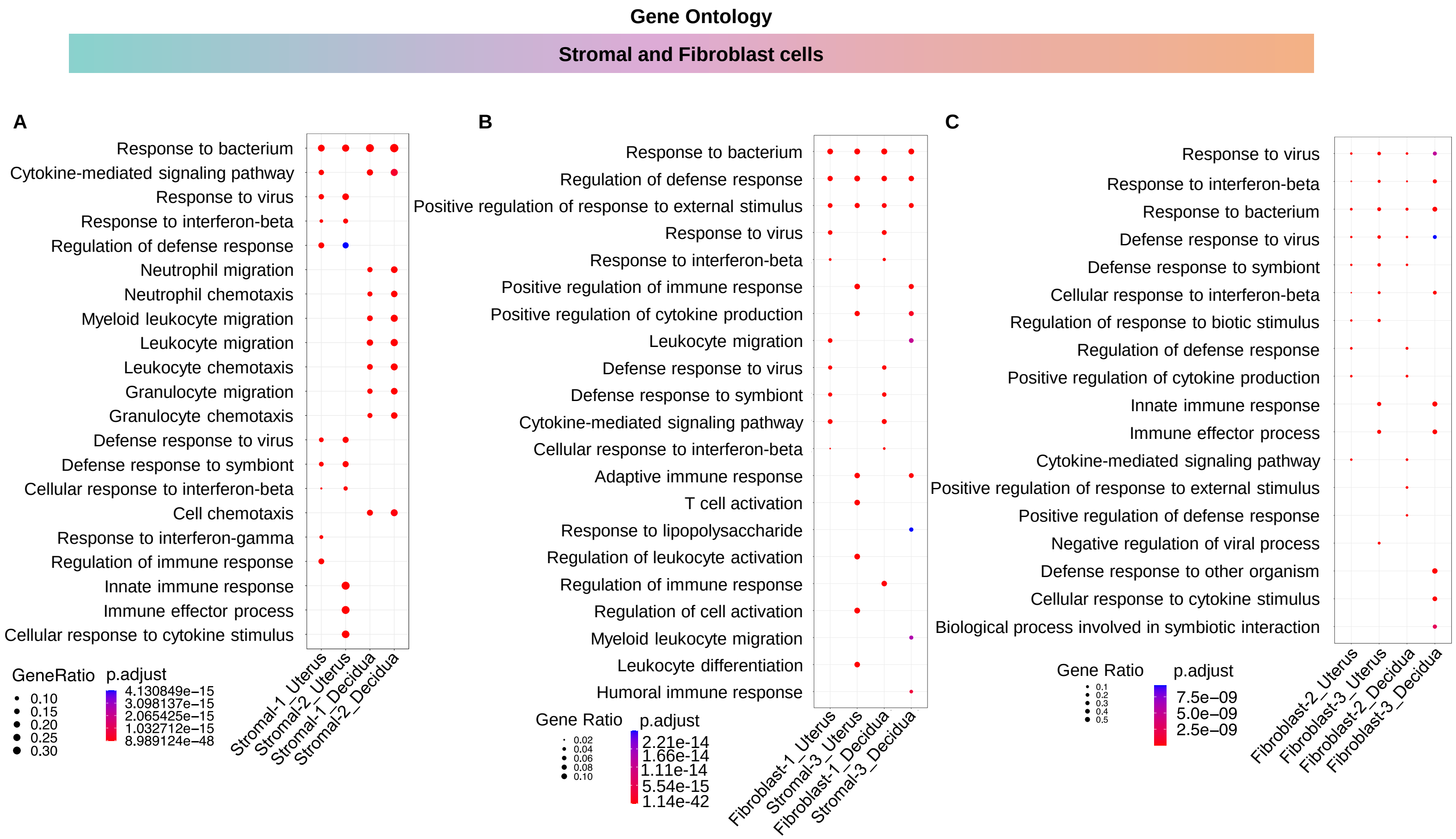


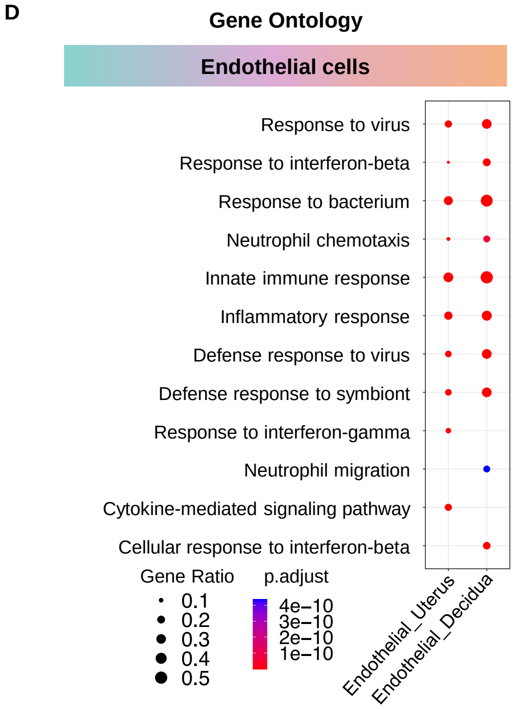


**Figure S4. The uterus and decidua share enrichment of biological processes in preterm labor, related to Figure 3.** Cluster profiler dot plots showing the Gene Ontology (GO) biological processes enriched with preterm labor in **(A)** Stromal-1 and Stromal-2, **(B)** Stromal-3 and Fibroblast-1, **(C)** Fibroblast-2 and Fibroblast-3, and **(D)** Endothelial cells in the uterus and decidua. The size and color of each dot represent gene ratio and significance level, respectively. A 1-sided Fisher's exact test was used.

### Uterus

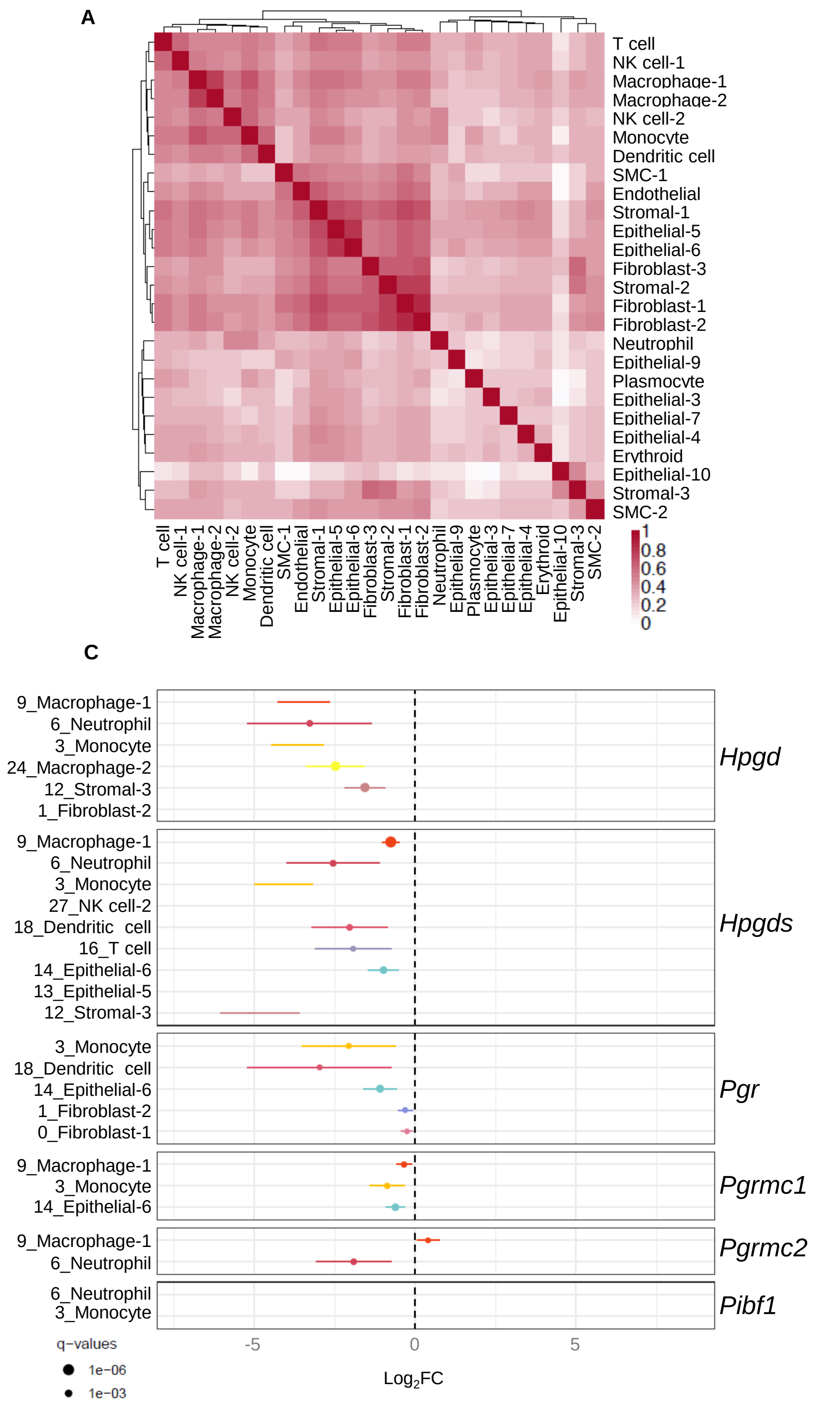


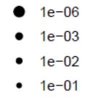


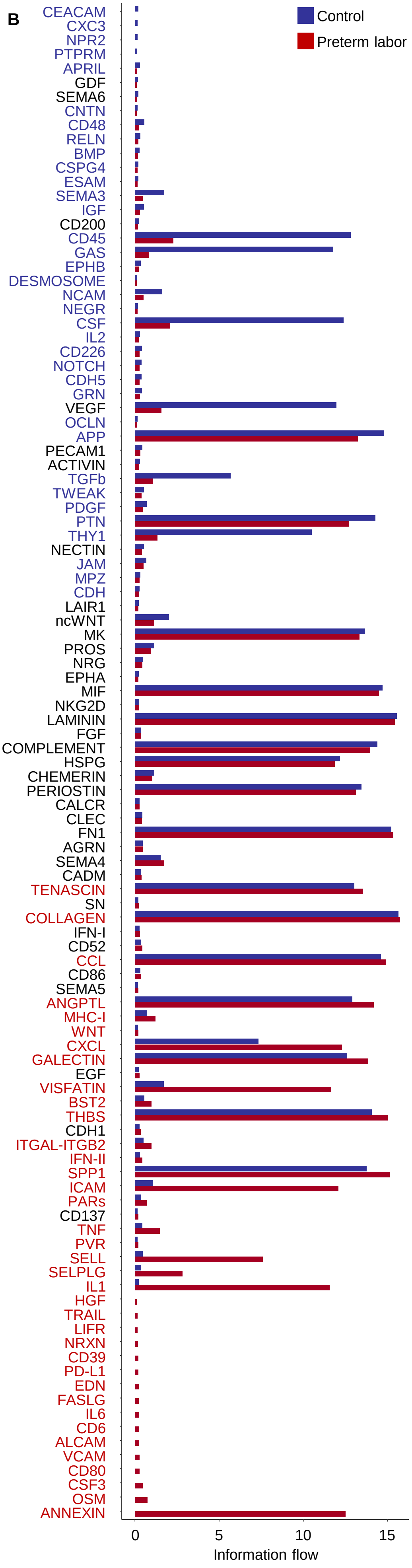


**Figure S5. Preterm labor-associated enrichment of signaling pathways in uterine cell types, related to Figure 4. (A)** Heatmap showing correlations among uterine cell types where red and white blocks signify increased and decreased correlation, respectively. Pearson correlation tests was used. **(B)** Bar plots showing the high expression of specific signaling pathways in the uterus of control mice (blue bars/pathway names) or mice with preterm labor (red bars/pathway names). Abbreviations used: SMC, smooth muscle cell; NK cell, natural killer cell. **(C)** Forest plot showing the log$_2$(FC, fold change) and 95% confidence intervals of differentially expressed (DEGs) across cell types in murine uterus. DEGs shown are significant with FDR (q < 0.1).

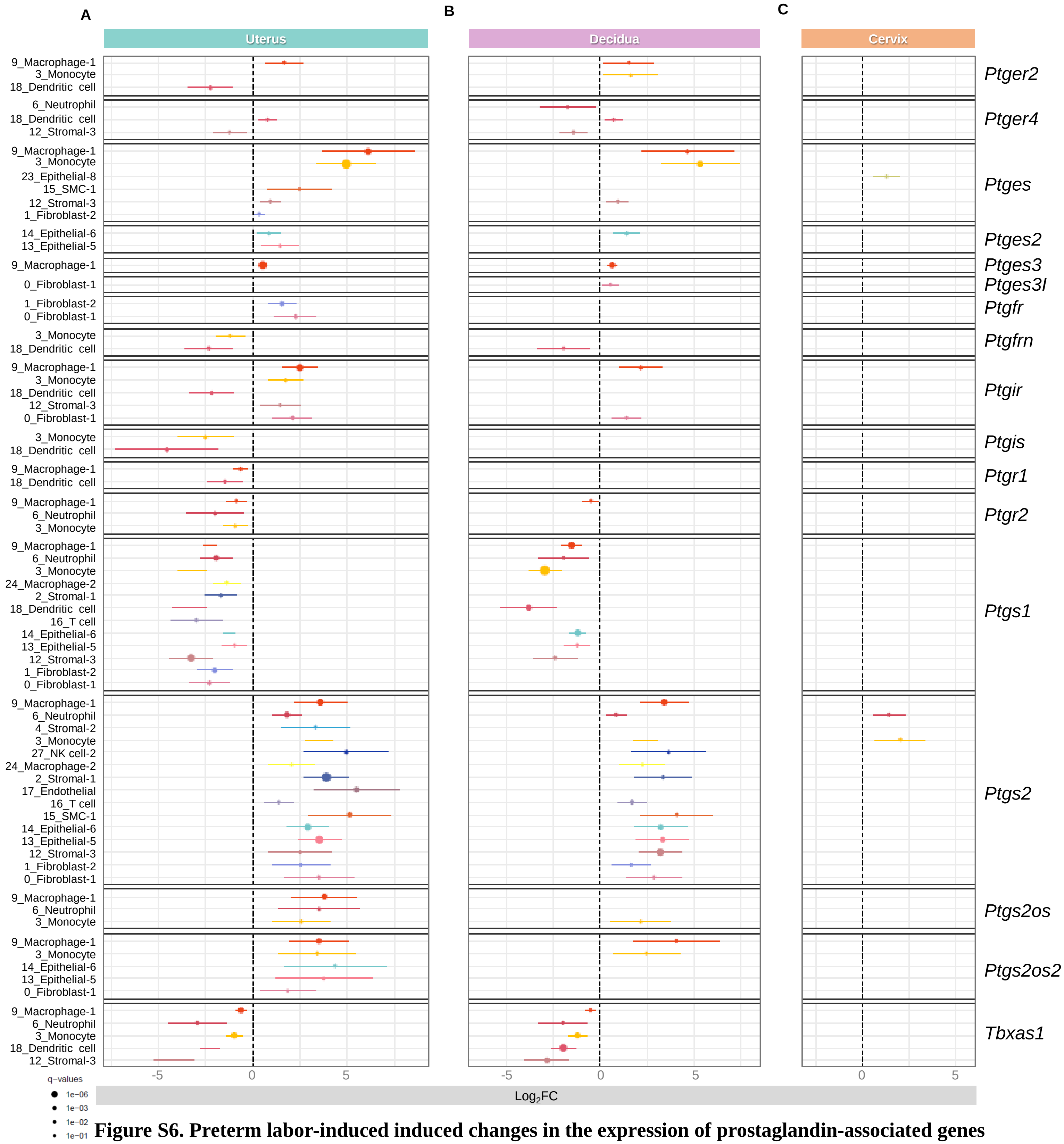


**Figure S6. Preterm labor-induced induced changes in the expression of prostaglandin-associated genes in different cell types across the reproductive tissues.** Forest plot showing the $\log_2$(FC, fold change) and 95% confidence intervals of differentially expressed (DEGs) in selected cell types across the murine **(A)** uterus, **(B)** decidua and **(C)** cervix. DEGs shown are significant with FDR (q < 0.01).

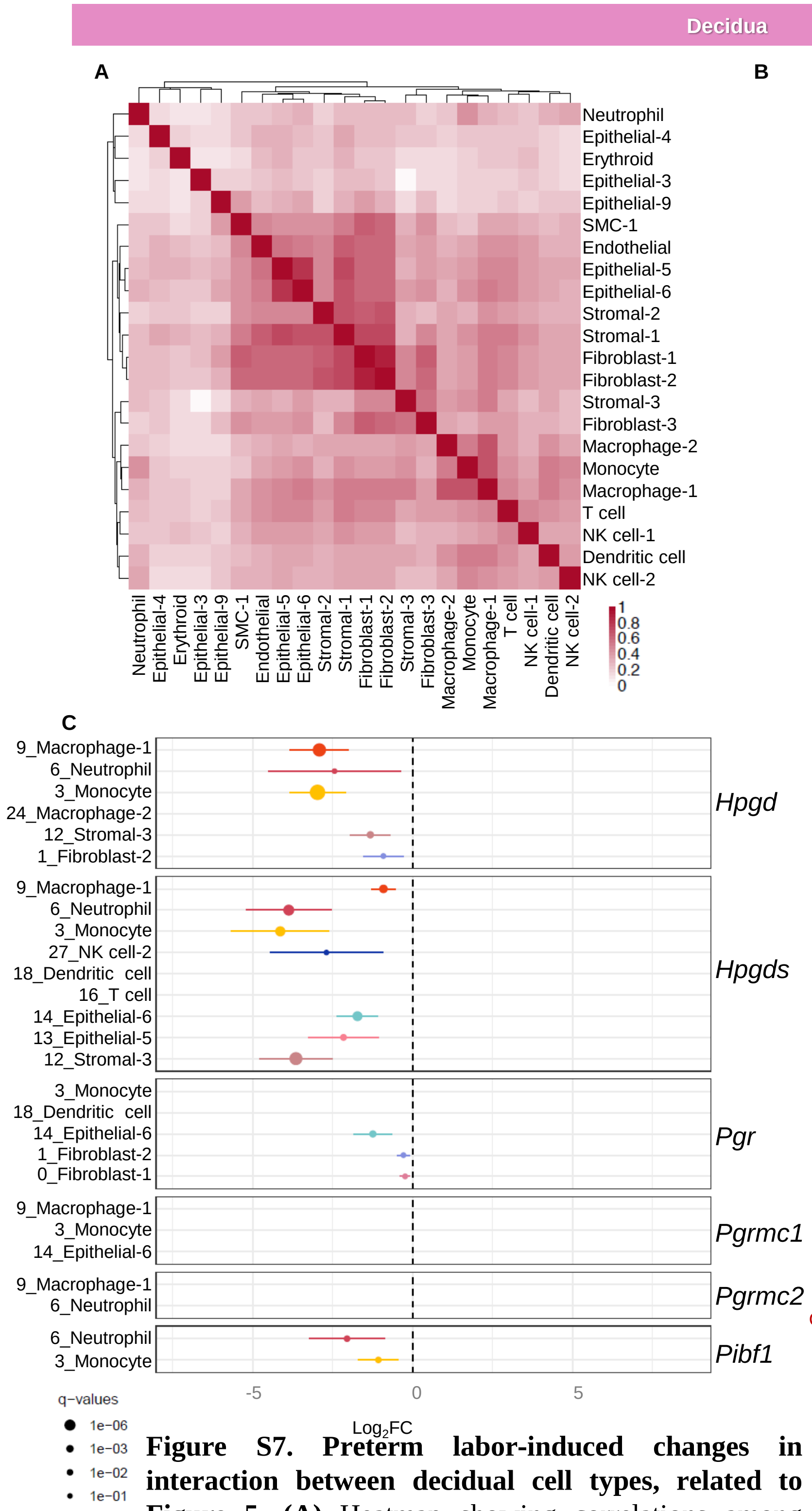


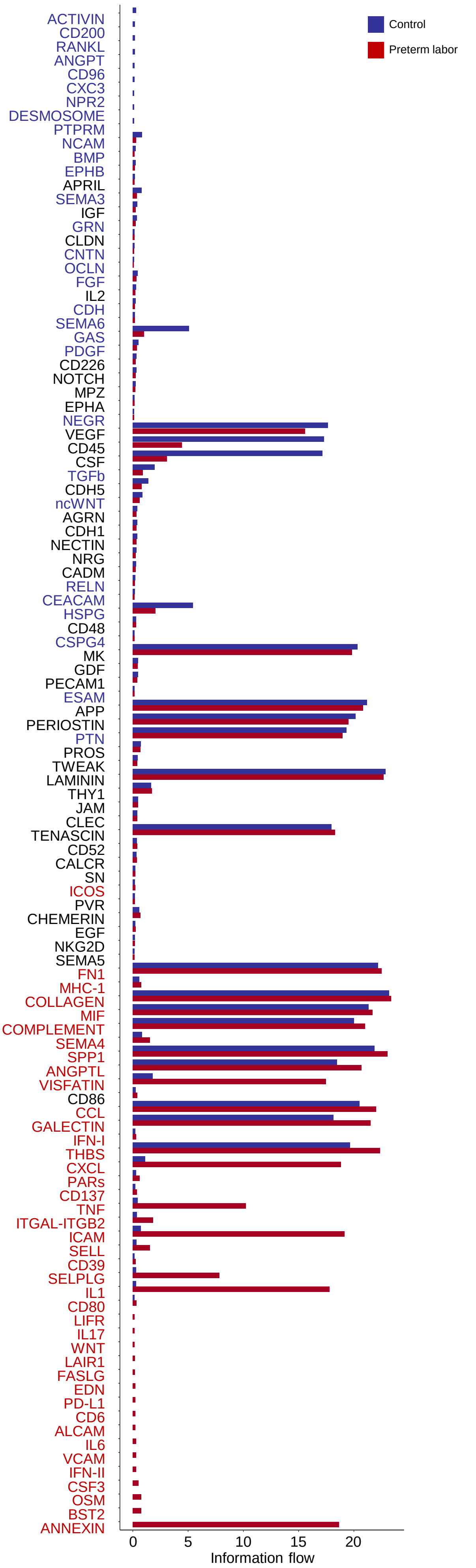


**Figure S7. Preterm labor-induced changes in interaction between decidual cell types, related to Figure 5. (A)** Heatmap showing correlations among decidual cell types where red and white blocks signify increased and decreased correlation, respectively. Pearson correlation tests was used. **(B)** Bar plots showing the high expression of specific signaling pathways in the decidua of control mice (blue bars/pathway names) or mice with preterm labor (red bars/pathway names). Abbreviations used: SMC, smooth muscle cell; NK cell, natural killer cell. **(C)** Forest plot showing the $\log_2$(FC, fold change) and 95% confidence intervals of differentially expressed (DEGs) across cell types in murine decidua. DEGs shown are significant with FDR ($q < 0.1$).

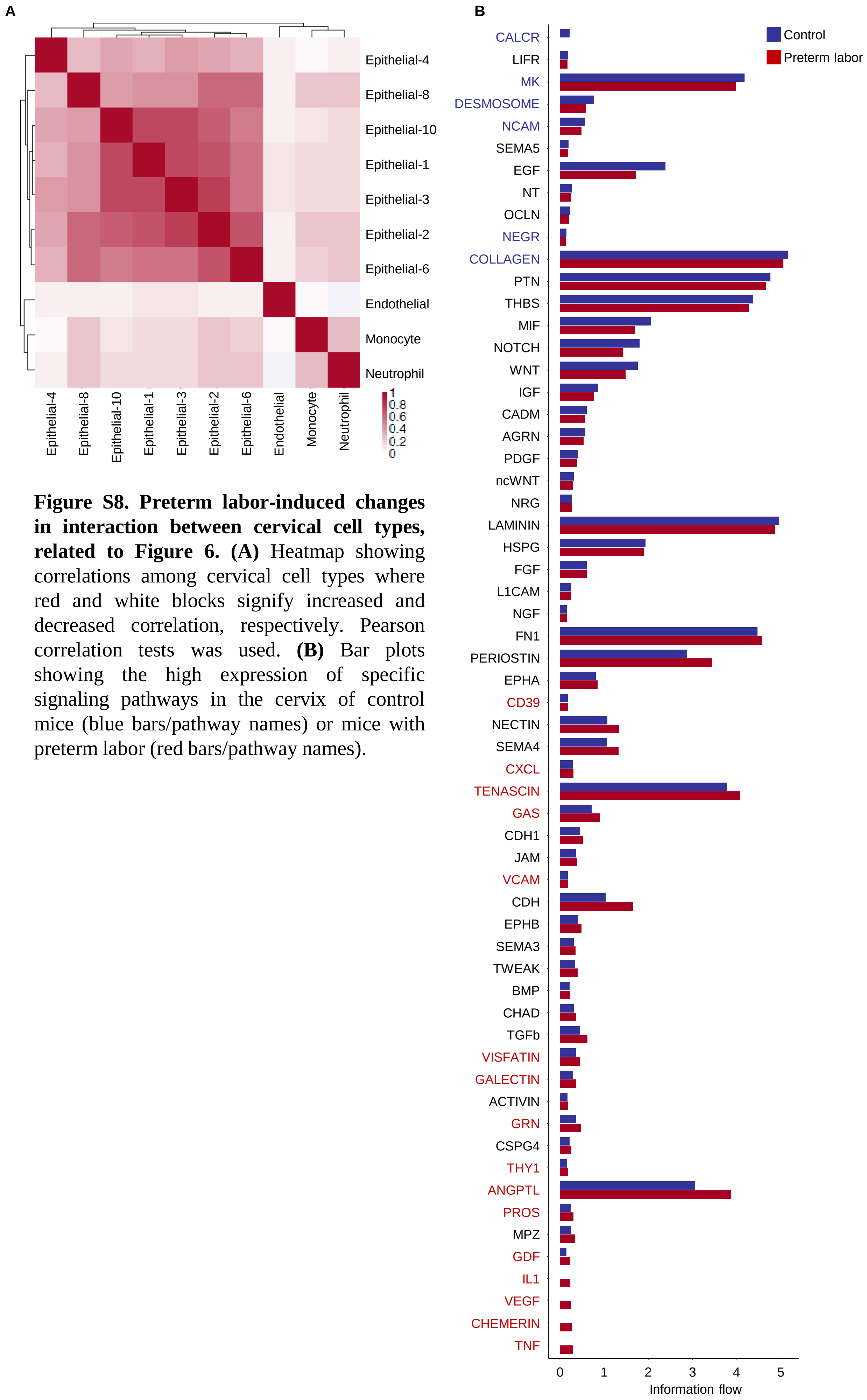


**Figure S8. Preterm labor-induced changes in interaction between cervical cell types, related to Figure 6. (A)** Heatmap showing correlations among cervical cell types where red and white blocks signify increased and decreased correlation, respectively. Pearson correlation tests was used. **(B)** Bar plots showing the high expression of specific signaling pathways in the cervix of control mice (blue bars/pathway names) or mice with preterm labor (red bars/pathway names).

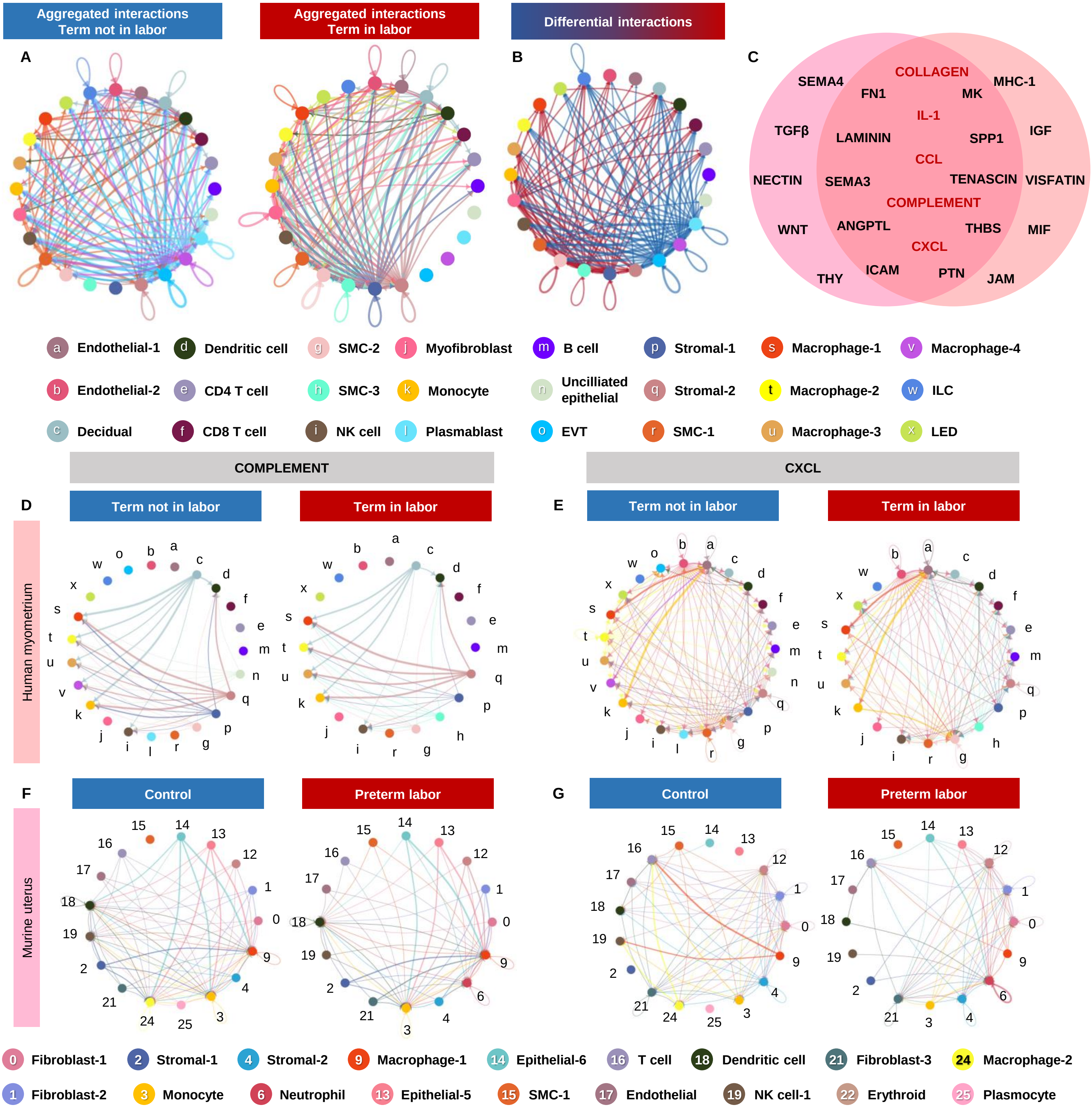


**Figure S9. Labor-associated signaling in the human myometrium partially overlaps with preterm labor-associated changes in the murine uterus, related to Figure 7. (A)** Circle plots showing the top aggregated interactions among cell types in the myometrium from humans without (left) or with labor at term (right). Each node represents a cell type and the interaction is shown by lines color-coded based on the sender cell. Representation of aggregated interactions with $p < 0.05$ using cell chat. **(B)** Circle plot showing the increased (red) or decreased (blue) signaling interactions in the human myometrium in labor compared to controls without labor. Representation of top 25% differential interaction strength. **(C)** Venn diagram showing the overlap in upregulated signaling pathways between the murine uterus in preterm labor (left circle, pink) and the human myometrium in term labor (right circle, orange). Shared labor- and inflammation-associated pathways are highlighted in red. **(D-E)** Circle plots representing the top 25% human myometrial cell-cell communications inferred for the Complement and CXCL pathways for the Term not in labor and Term in labor groups. **(F-G)** Circle plots representing the top 25% murine uterine cell-cell communications inferred for the Complement and CXCL pathways for the control and preterm labor groups. Abbreviations used: SMC, smooth muscle cell; NK cell, natural killer cell; EVT, extravillous trophoblast; ILC, innate lymphoid cell; LED, lymphoid endothelial decidual cell.
