## Supplementary material for "The single-cell atlas of the murine reproductive tissues during preterm labor": Key Resources Table

| REAGENT or RESOURCE | SOURCE | IDENTIFIER |
| --- | --- | --- |
| Antibodies | | |
| the Monoclonal Rabbit Anti-Mouse CD45 | Cell Signaling Technology | Cat#70257S;  RRID: AB_2799780 |
| Rabbit FLEX Universal Negative Control | Agilent | Cat# IR60066-2 |
| Monoclonal Rabbit Anti-Mouse F4/80 | Cell Signaling Technology | Cat# 70076S; RRID:AB_2799771 |
| Monoclonal Rabbit Anti-Mouse CD3ε | Cell Signaling Technology | Cat# 78588S; RRID:AB_2889902 |
| Monoclonal Rabbit Anti-Mouse Klrb1c/CD161c | Cell Signaling Technology | Cat# 39197S; RRID:AB_2892989 |
| Polyclonal Rabbit Anti-Mouse Ly6C | HuaBio | Cat# HA500088 |
| Monoclonal Rabbit Anti-Mouse Ly6G | Cell Signaling Technology | Cat# 87048S; RRID:AB_2909808 |
| Bacterial and virus strains | | |
| Escherichia coli | ATCC | ATCC 12014 |
| Critical commercial assays | | |
| Umbilical Cord Dissociation Kit, human | Miltenyi Biotec | Cat# 130-105-737 |
| Dead Cell Removal Kit | Miltenyi Biotec | Cat# 130-090-101 |
| Chromium Next GEM Single Cell 3’ GEM, Library &Gel beads Kit | 10x Genomics | PN:1000121 |
| Chromium Next GEM Chip G Single Cell Kit | 10x Genomics | PN:1000120 |
| Single Index Kit T Set A | 10x Genomics | PN: 1000213 |
| SPRIselect Reagent | Beckman Coulter | Item Number:  B23318 |
| Deposited data | | |
| scRNA-sequencing | This paper | GSE200289 |
| Experimental models: Organisms/strains | | |
| Mouse: C57BL/6 | The Jackson Laboratory | RRID:IMSR_JAX:000664 |
| Mouse: BALB/cByJ | The Jackson Laboratory | RRID:IMSR_JAX:001026 |
| Software and algorithms | | |
| Cell Ranger version 4.0.0 | 10x Genomics | http://www.10xgenomics.com |
| STAR aligner | Dobin et al., 2013 | https://github.com/alexdobin/STAR |
| Demuxlet | Kang et al., 2018 | https://github.com/statgen/demuxlet |
| SoupX version 1.5.2 | Young et al., 2020 | https://github.com/constantAmateur/Soup |
| DoubletFinder 2.0.3 | McGinnis., 2019 |  |
| Seurat version 4.0.3 | Hafemeister et al., 2019; Stuart et al., 2019 | https://satijalab.org/seurat/ |
| Harmony package in R version 1.0.0 (R package from CRAN) | Korsunsky et al., 2019 | N/A |
| SingleR package in R version 1.6.1 | Aran et al., 2019 | https://doi.org/doi:10.18129/B9.bioc.SingleR |
| Mouse Cell Atlas and single-cell MCA (scMCA) package in R version 0.2.0 | Han et al., 2018 | http://bis.zju.edu.cn/MCA/search.html  https://github.com/ggjlab/scMCA |
| DESeq2 R package version 1.32.0 | Love et al., 2014 | https://doi.org/doi:10.18129/B9.bioc.DESeq2 |
| clusterProfiler in R version 4.0.4 | Yu et al., 2012 | https://doi.org/doi:10.18129/B9.bioc.clusterProfiler |
| CellChat version 1.1.2 (R package from CRAN) | Jin et al., 2021 | N/A |
| ggalluvial version 0.12.3 (R package from CRAN) |  | N/A |
| ggplot2 version 3.3.5 (R package from CRAN) |  | N/A |
| Uniform Manifold Approximation and Projection for Dimension Reduction (UMAP) algorithm | McInnes et al., 2018;  Becht et al., 2018 | N/A |
| Other Data sets | | |
| Human uterine scRNA-seq data | Pique-Regi et al., 2022 | Accession number  phs001886.v4.pl |
